# Mechanoadaptive focal adhesion–actomyosin coupling organizes diverse modes across ECM stiffness

**DOI:** 10.64898/2026.10.26.762083

**Authors:** Eiji Matsumoto, Shinji Deguchi

**Affiliations:** Graduate School of Engineering Science, The University of Osaka, 1-3 Machikane-yama, Toyonaka, Osaka 560-8531, Japan; Global Center for Medical Engineering and Informatics, The University of Osaka, Osaka, Japan; R^3^ Institute for Newly-Emerging Science Design, The University of Osaka, Osaka, Japan

**Keywords:** Mechanoadaptation, Focal adhesion, ECM stiffness, Mechanical homeostasis, Mechanical memory, Multistability

## Abstract

Focal adhesions mechanically couple the actomyosin cytoskeleton to the extracellular matrix (ECM), contributing to mechanosensing and mechanical homeostasis. They span diverse states in response to the mechanical environment, from nascent adhesions and focal complexes to mature and supermature adhesions, undergoing dynamic remodeling. However, how ECM-dependent force generation and internal mechanoadaptation jointly shape the emergence and stability of these states remains unclear. Here, we develop a four-variable dynamical model comprising adhesion state, transmitted force, reinforcement activity, and an adaptive mechanical stress target. As ECM stiffness varies, the model generates distinct modes while regulating mechanical stress toward this target. The force-response timescale further modifies the stability structure, giving rise to a stable oscillatory mode. Within multistable regimes, adhesion-mode selection depends not only on current ECM stiffness but also on its history. These results show that diverse adhesion modes can emerge from a common mechanoadaptive system, with ECM stiffness history influencing mode selection.

**GRAPHICAL ABSTRACT:** 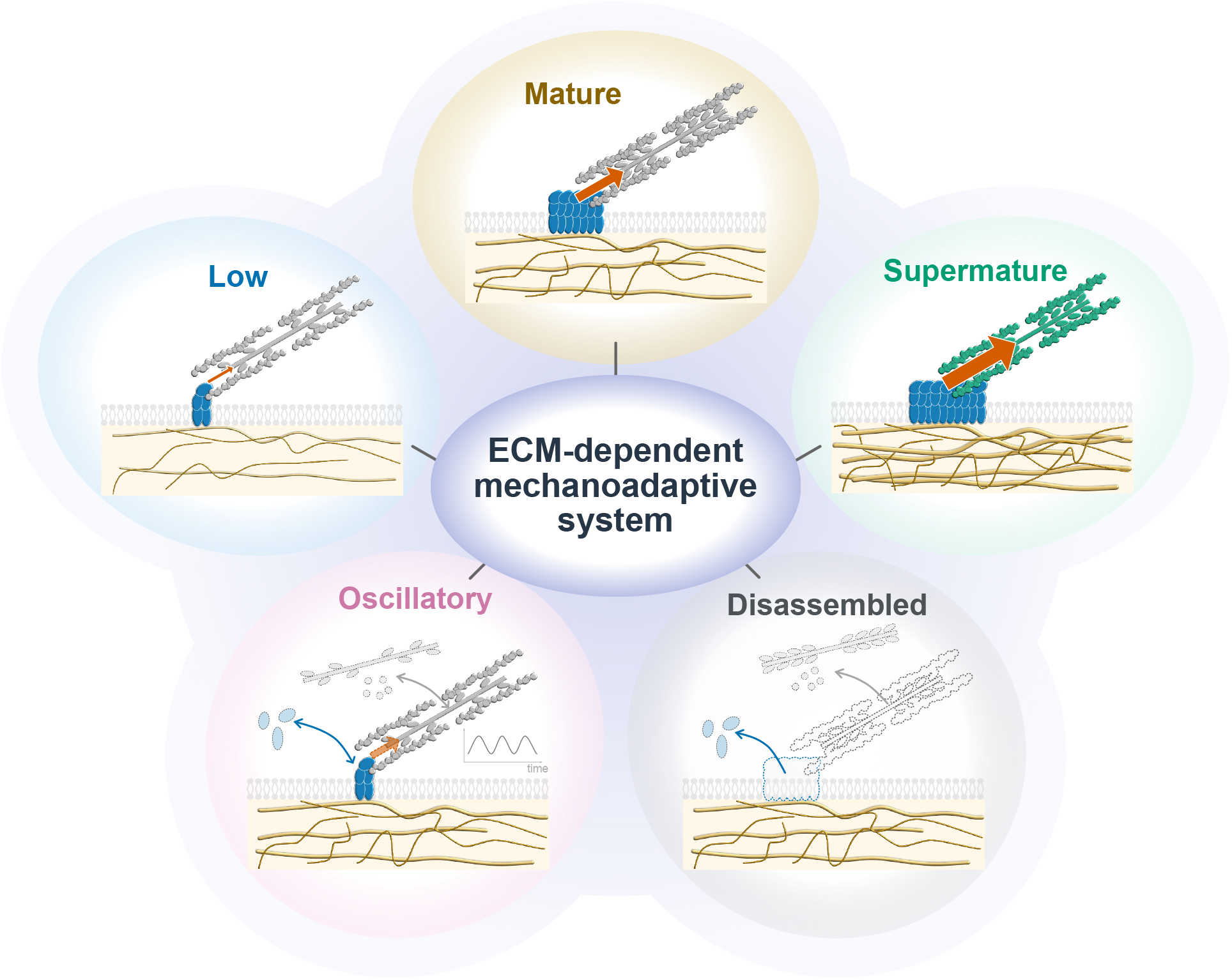

## I. INTRODUCTION

Focal adhesions are mechanosensitive cell–matrix interfaces that couple the actin cytoskeleton to the extra-cellular matrix (ECM), enabling force transmission and mechanotransduction [1–3]. Actomyosin-generated tension promotes focal adhesion growth and maturation [4– 7]. ECM stiffness, in turn, regulates cellular contractility through focal-adhesion mechanosensing and RhoA– ROCK–myosin II signaling [8–10]. Such mechanosensitive interactions also contribute to mechanical homeostasis (or tensional homeostasis), in which stress sustained by focal adhesions is regulated toward characteristic levels despite changes in the mechanical environment [11, 12].

Focal adhesions exhibit diverse states that differ in organization and force transmission. Their development and maturation involve changes in molecular organization and signaling, along with stronger coupling to actomyosin stress fibers [6, 13–17]. In mature adhesions, transmitted force generally scales with adhesion area, suggesting a characteristic traction stress consistent with mechanical homeostasis [5, 18, 19]. Incorporation of *α*-smooth muscle actin (*α*-SMA) into stress fibers enhances cellular traction and promotes further reinforcement, with supermature adhesions sustaining higher stress [19– 22]. Focal adhesions also undergo turnover through assembly and disassembly [23], exhibiting traction dynamics ranging from relatively stable levels to fluctuations and oscillations [24–26].

Theoretical frameworks have addressed focal adhesion dynamics from multiple perspectives. Force-dependent adhesion models examined cluster stability and bistability [27–29], force regulation by nascent adhesions [30], and growth, maturation, and dynamic transitions [31– 34]. Motor–clutch models described stiffness-dependent force transmission and load-and-fail dynamics [35, 36], with extensions incorporating reinforcement [37], multiscale mechanics [38], oscillations [39], and migration modes [40]. Models coupling cytoskeletal mechanics with adhesion regulation captured stiffness-dependent hysteresis [41], load-dependent contractility [42], and oscillatory behaviors [25, 43], while other models identified traction bistability [44] and multiple homeostatic targets [45]. However, how ECM-dependent force generation and adaptive stress regulation determine the existence and stability of diverse adhesion modes remains unclear.

Here, we develop a four-variable dynamical model of focal adhesion–actomyosin coupling comprising adhesion state, transmitted force, reinforcement activity, and an adaptive mechanical stress target. The adaptive target sets a state-dependent reference level, and ECM stiffness regulates force generation. We characterize how ECM stiffness and the force-response timescale organize Disassembled, Low, Mature, Supermature, and Oscillatory modes, including multistable regimes. We further examine how ECM stiffness history influences adhesion-mode selection under the same final stiffness.

## II. MECHANOADAPTIVE MODEL OF FOCAL ADHESION–ACTOMYOSIN COUPLING

### A. Model setup

The model represents a focal adhesion–actomyosin system coupled to ECM stiffness with four state variables: adhesion state *A* (represented by focal adhesion area); transmitted actomyosin force *F* ; reinforcement activity *R*; and an adaptive mechanical stress target *S* (Fig. 1A). ECM stiffness *E* serves as an external mechanical input, and *F/A* denotes the average stress sustained by the adhesion (Fig. 1B). The feedback architecture is motivated by experimentally observed relationships involving force-dependent remodeling and force–area coupling [4–6], ECM-sensitive force generation [8–10], and reinforcement-dependent contractility [19, 46, 47]. State-dependent stress levels and mechanical homeostasis motivate *S* as an adaptive stress target [5, 11, 12, 19, 45].

**FIG. 1.**
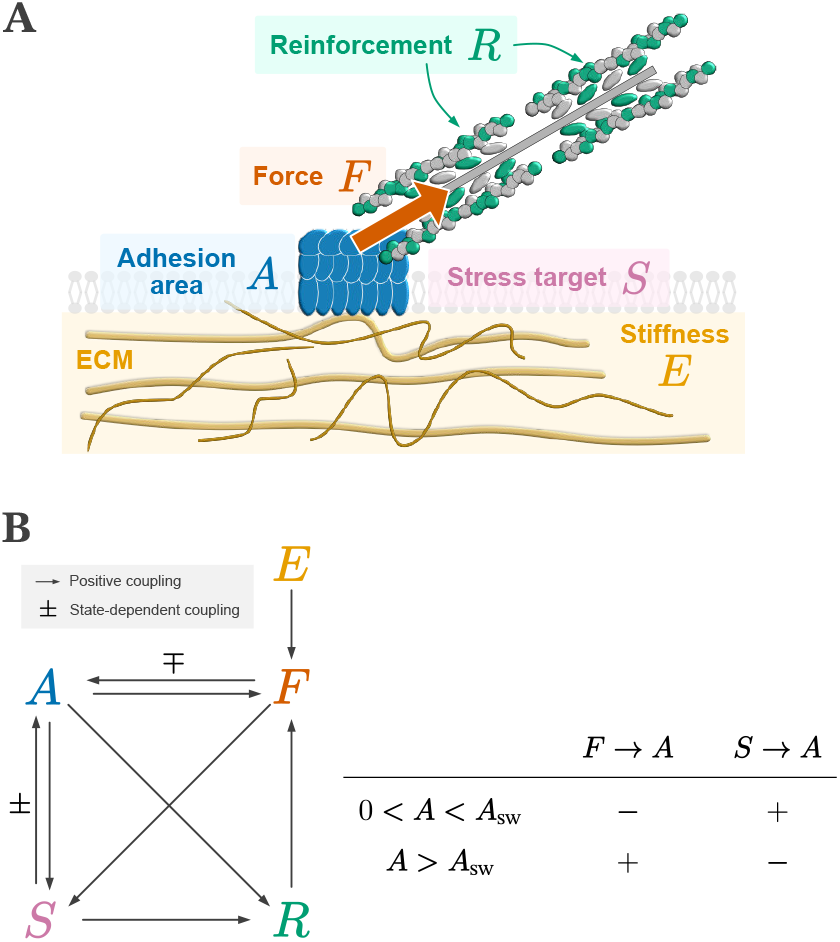
State-dependent mechanoadaptive feedback couples ECM stiffness to focal adhesion–actomyosin regulation. (A) Schematic of the cell–ECM adhesion system showing ECM stiffness *E* and four state variables: adhesion state *A*, transmitted force *F*, reinforcement activity *R*, and adaptive mechanical stress target *S*. (B) State-dependent feedback interactions in the model.

### B. Model equations

Saturating regulatory responses are represented by the Hill function

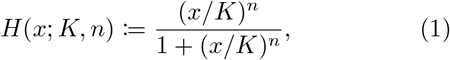

where *K* is the half-saturation value and *n* controls the response nonlinearity. The model is governed by

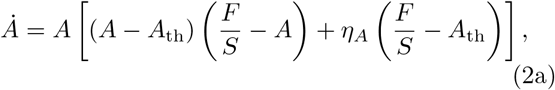

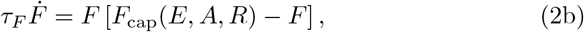

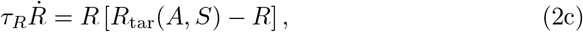

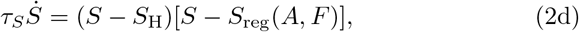

with

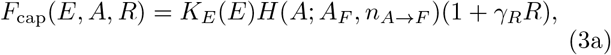

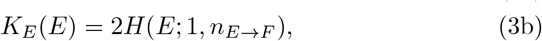

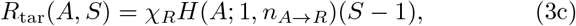

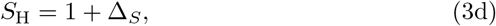

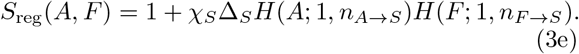

Equation (2a) uses a multiplicative form in which adhesion remodeling scales with *A* and responds to the force–adhesion mismatch *F/S* − *A* relative to the stress target. The factor *A* − *A*_th_ introduces a maturation-dependent switch in mechanical feedback, reflecting the shift from early adhesion behavior to tension-dependent focal-adhesion growth [7, 14, 48, 49]. A small offset *η*_*A*_ allows *A* to cross the threshold *A* = *A*_th_.

In Eq. (2b), we adopt logistic-type dynamics so that positive transmitted force approaches the state-dependent capacity *F*_cap_(*E, A, R*) when *E, A*, and *R* are fixed. The capacity *F*_cap_ is positively regulated by ECM stiffness [8, 9], adhesion-dependent actomyosin contractility [10, 50], and reinforcement [19, 46]. In Eq. (2c), reinforcement adjusts toward *R*_tar_, representing an adhesion-dependent response to an elevated stress target [19, 51, 52]. In Eq. (2d), the stress target has two stationary branches: the fixed high target *S*_H_ and the mechanically regulated target *S*_reg_(*A, F* ). As *S*_reg_ varies with *A* and *F*, the stress target can switch between the two branches.

Here, *A*_th_ denotes the adhesion threshold and *A*_*F*_ the half-saturation adhesion state for the *A* → *F* coupling. The gains *γ*_*R*_, *χ*_*R*_, and *χ*_*S*_ control reinforcement-dependent force amplification, the reinforcement target, and mechanically regulated stress-target elevation, respectively, while Δ_*S*_ sets the separation between the baseline and high-stress targets. The subscripts of *n*_*X→Y*_ identify the corresponding Hill couplings, and *τ*_*F*_, *τ*_*R*_, and *τ*_*S*_ set the response timescales of force, reinforcement, and stress-target adaptation relative to adhesion remodeling. We consider *A* ≥ 0, *F* ≥ 0, *R* ≥ 0, *E* ≥ 0, and the physical stress-target range 1 ≤ *S* ≤ 1 + Δ_*S*_. All variables are nondimensional, with unit reference levels determined by the chosen scaling; the corresponding dimensional formulation and nondimensionalization are given in Supplementary Text S1.

### C. State-dependent feedback

The state dependence of mechanical feedback follows from Eq. (2a):

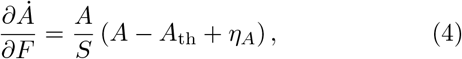

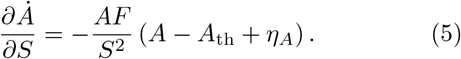

The switching threshold is

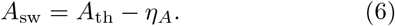

Since *η*_*A*_ ≪ *A*_th_, *A*_sw_ ≃ *A*_th_. For *A < A*_sw_, the force-to-adhesion coupling is negative and the stress-target-to-adhesion coupling is positive, whereas both signs reverse for *A > A*_sw_ (Fig. 1B).

### D. Parameterization

We use a reference parameter set throughout the analysis unless otherwise stated (Table I). The offset *η*_*A*_ = 10^*−*4^ is much smaller than *A*_th_ = 0.1, keeping the switching threshold *A*_sw_ close to *A*_th_. We vary the force-response timescale *τ*_*F*_ and ECM stiffness *E* to examine their effects on adhesion dynamics. The main stationary and oscillatory features persist under the parameter variations examined in Figs. S2 and S3.

**TABLE I.** Reference parameters of the dimensionless model.

| Parameter | Value | Role |
| --- | --- | --- |
| $A_{th}$ | 0.1 | Adhesion-state threshold |
| $\eta_A$ | $10^{-4}$ | Adhesion-response offset |
| $A_F$ | 0.5 | Adhesion-to-force half-saturation |
| $\gamma_R$ | 3 | Reinforcement-to-force gain |
| $\chi_R$ | 0.5 | Reinforcement gain |
| $\chi_S$ | 2 | Stress-target regulation gain |
| $\Delta_S$ | 1 | Stress-target range |
| $n_{E \rightarrow F}$ | 1.2 | ECM-to-force nonlinearity |
| $n_{A \rightarrow F}$ | 1.2 | Adhesion-to-force nonlinearity |
| $n_{A \rightarrow R}$ | 2 | Adhesion-to-reinforcement nonlinearity |
| $n_{A \rightarrow S}$ | 2 | Adhesion-to-target nonlinearity |
| $n_{F \rightarrow S}$ | 2 | Force-to-target nonlinearity |
| $\tau_F$ | 1 | Relative force response time |
| $\tau_R$ | 1 | Relative reinforcement response time |
| $\tau_S$ | 1 | Relative stress-target response time |

## III. RESULTS

### A. ECM stiffness organizes multiple stationary adhesion states

We first examined how ECM stiffness organizes the stationary states of the mechanoadaptive system. The Disassembled stationary state (*A, F, R, S*) = (0, 0, 0, 1) is locally stable for any ECM stiffness, representing the absence of a force-bearing adhesion. Multiple positive equilibrium branches were also identified (Fig. 2A–D). The stable low- and high-adhesion branches overlapped over a finite range of ECM stiffness, allowing stationary coexistence. Along the stable high-adhesion branches, adhesion state and transmitted force generally increased with ECM stiffness, and equilibria with larger adhesion states generally exhibited greater reinforcement activity. On the regulated high-adhesion branch, the adaptive target *S* increased toward *S*_H_ = 2 and reached the fixed-high branch *S* = *S*_H_ at higher stiffness. In these equilibria, the adhesion stress *F/A* matched the adaptive target *S* (maximum relative mismatch *<* 0.08%; Supplementary Text S2), providing a stationary-state basis for mechanical homeostasis.

**FIG. 2.**
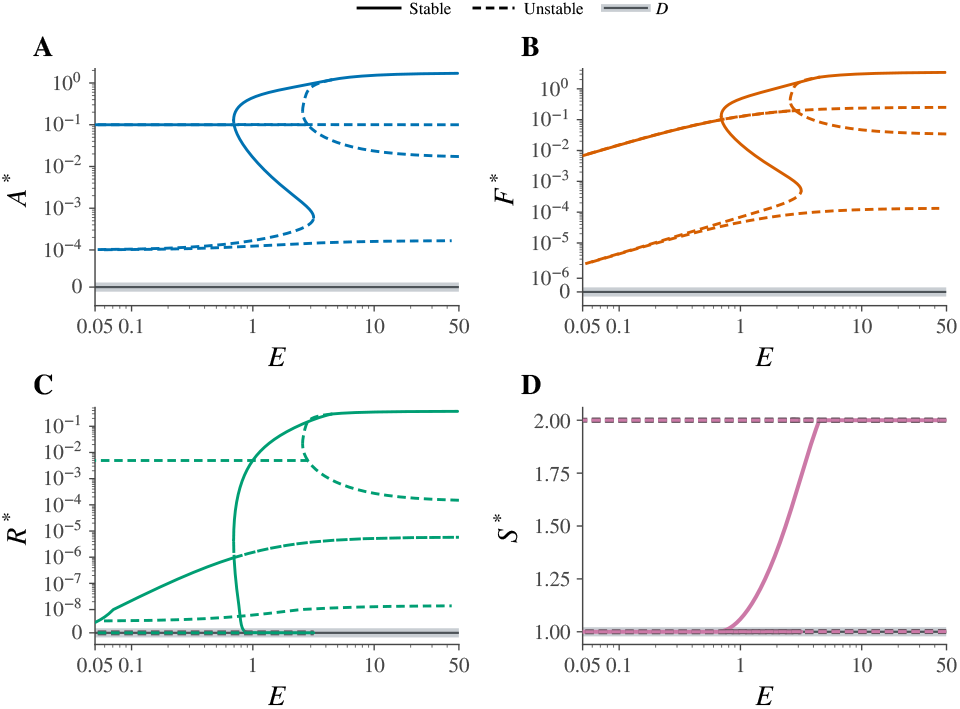
ECM stiffness organizes multiple stationary adhesion states. Stationary branches are shown as functions of ECM stiffness *E*. (A–D) Stationary values of *A, F, R*, and *S*, respectively. Solid and dashed lines denote stable and unstable positive equilibria, and the Disassembled state is shown in gray.

### B. Diverse adhesion modes emerge across ECM stiffness and the force-response timescale

We next examined how ECM stiffness and the force-response timescale jointly organize dynamical adhesion modes (Fig. 3A). These modes are defined in Table II. Consistent with Fig. 2, the stationary modes were organized from Low to Mature and then to Supermature with increasing *E*, with Low–Mature coexistence over an intermediate range. At intermediate *E*, increasing *τ*_*F*_ destabilized the Low equilibrium through a Hopf bifurcation. At *E* = 1, the Low equilibrium underwent a supercritical Hopf bifurcation at *τ*_*F*_ ≃ 11.91 (Fig. 3B), and the resulting periodic branch remained orbitally stable until the limit point of cycles (LPC) at *τ*_*F*_ ≃ 13.11 (Fig. 3C). The Mature equilibrium remained locally stable over the same interval, establishing Mature–Oscillatory coexistence.

**TABLE II.**
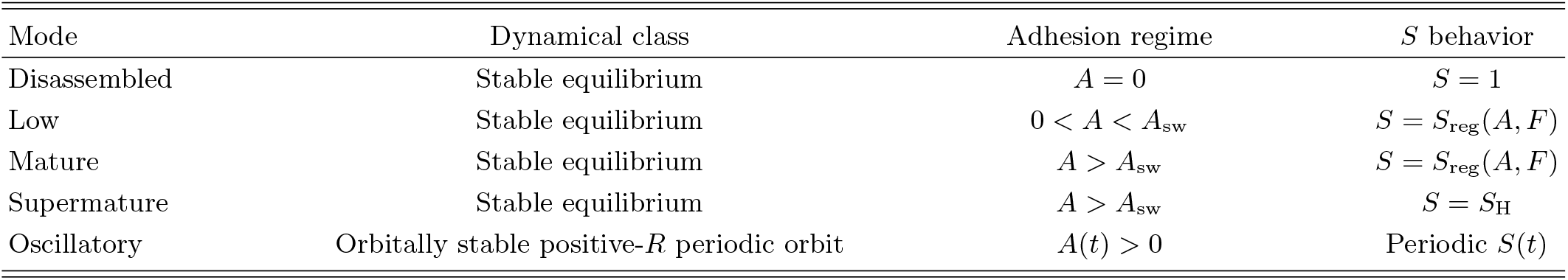
Definition of adhesion modes in the mechanoadaptive model. Disassembled denotes the stationary state (*A, F, R, S*) = (0, 0, 0, 1). Here, *A*_sw_ = *A*_th_ *− η*_*A*_ and *S*_H_ = 1 + Δ_*S*_ = 2.

| Mode | Dynamical class | Adhesion regime | $S$ behavior |
| --- | --- | --- | --- |
| Disassembled | Stable equilibrium | $A = 0$ | $S = 1$ |
| Low | Stable equilibrium | $0 < A < A_{sw}$ | $S = S_{reg}(A, F)$ |
| Mature | Stable equilibrium | $A > A_{sw}$ | $S = S_{reg}(A, F)$ |
| Supermature | Stable equilibrium | $A > A_{sw}$ | $S = S_H$ |
| Oscillatory | Orbitally stable positive-$R$ periodic orbit | $A(t) > 0$ | Periodic $S(t)$ |

**FIG. 3.**
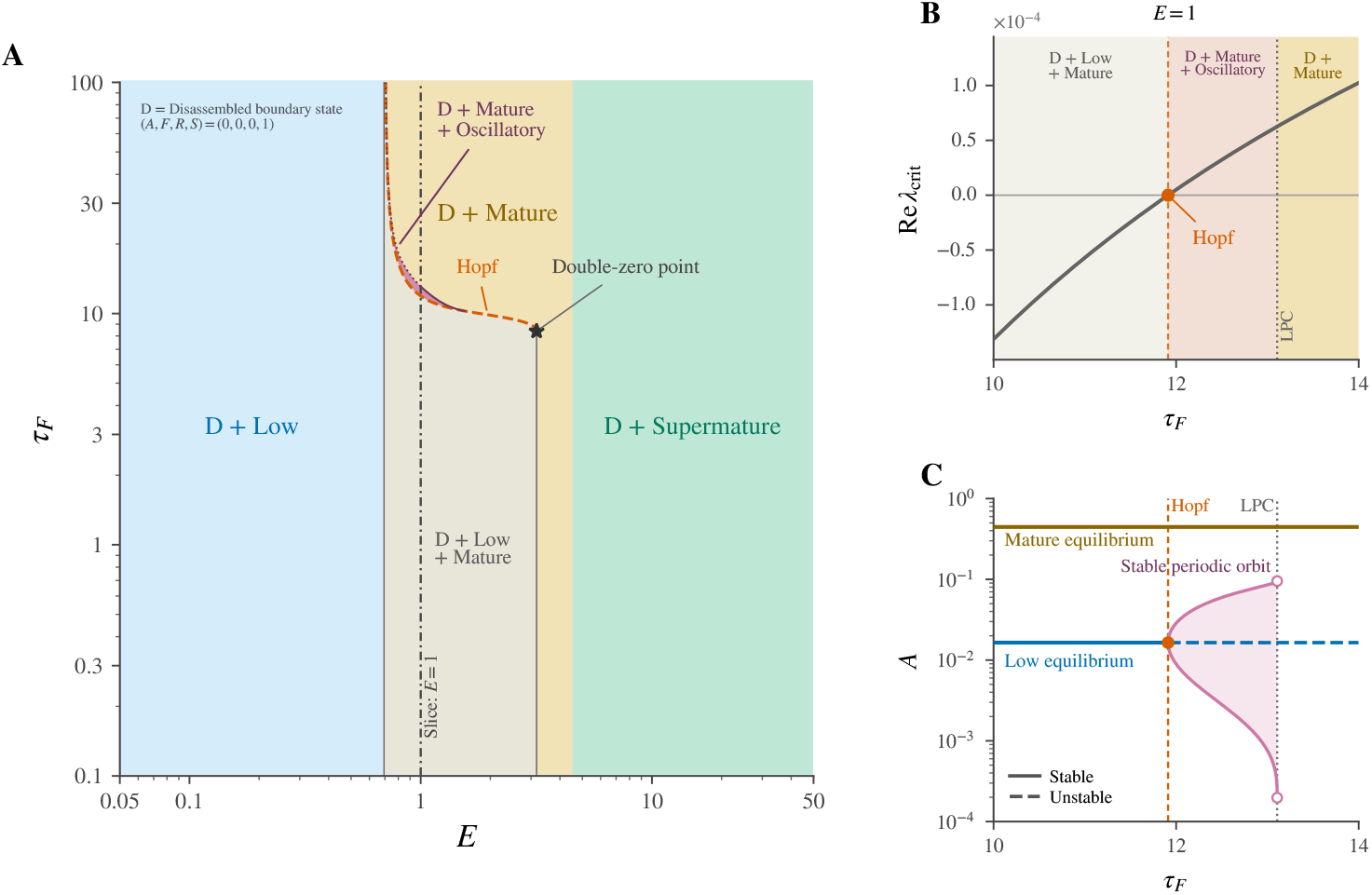
Distinct dynamical adhesion modes emerge across ECM stiffness and the force-response timescale. (A) Adhesion-mode organization in the (*E, τ*_*F*_ ) plane, showing the Hopf bifurcation curve and the outer extent of the positive-*R* periodic branch. Solid portions of the outer boundary indicate sections verified as limit points of cycles (LPCs), whereas the dotted portion marks the numerically verified extent of the positive-*R* periodic branch. The high-*E* boundary is shown through the last verified LPC section. The *E* = 1 slice analyzed in (B, C) and the spectral double-zero point are indicated. (B) Real part of the critical eigenvalue of the Low equilibrium, showing the Hopf crossing. (C) Adhesion-state values of the Low and Mature equilibria and the minimum and maximum *A* values of the positive-*R* periodic branch.

Representative trajectories approached the Disassembled, Low, Mature, and Supermature equilibria at distinct adhesion and force levels (Fig. 4A–D). The Oscillatory mode exhibited sustained oscillations along an orbitally stable periodic orbit (Fig. 4E). At *E* = 1, Low– Mature coexistence occurred at *τ*_*F*_ = 10 (Fig. 4F), and Mature–Oscillatory coexistence at *τ*_*F*_ = 12.5 (Fig. 4G). Different initial internal states led to distinct attractors under identical parameter conditions.

**FIG. 4.**
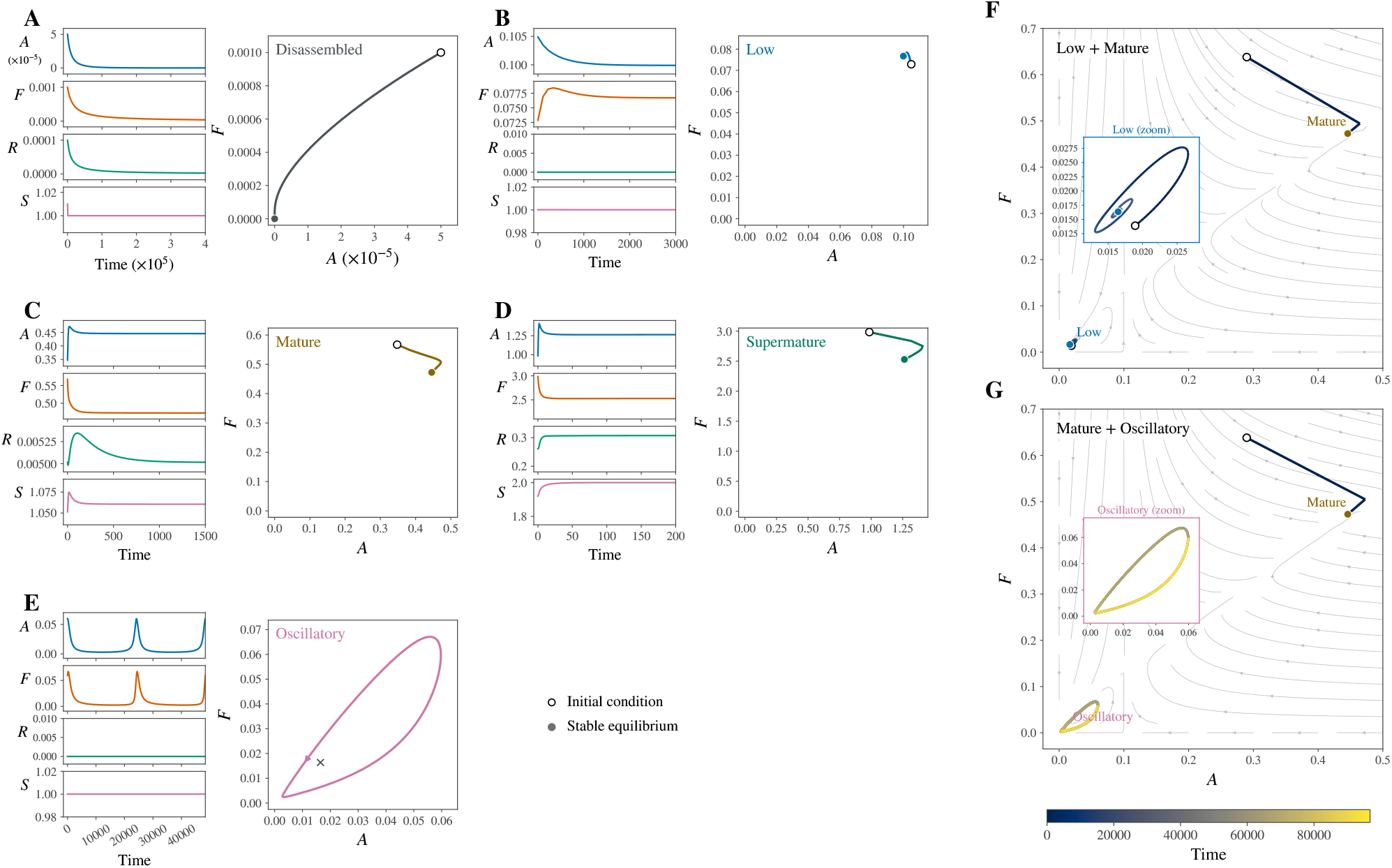
Representative adhesion dynamics and multistability in the mechanoadaptive system. (A–E) Time series and (*A, F* ) projections of the Disassembled, Low, Mature, Supermature, and Oscillatory modes. A gray cross marks the unstable Low equilibrium in (E). The parameter pairs (*E, τ*_*F*_ ) are (0.8, 16), (0.5, 10), (1, 10), (5, 10), and (1, 12.5) for (A)– (E), respectively, with *τ*_*R*_ = *τ*_*S*_ = 1. (F, G) Multistable dynamics projected onto the (*A, F* ) plane at *E* = 1: Low–Mature coexistence at *τ*_*F*_ = 10 (F) and Mature–Oscillatory coexistence at *τ*_*F*_ = 12.5 (G). Gray streamlines indicate local flow directions of the conditional 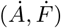 field evaluated at fixed *R* and *S*. Insets enlarge the Low and Oscillatory attractors in (F) and (G), respectively.

### C. ECM history shapes adhesion-mode selection

To examine the role of ECM stiffness history in adhesion-mode selection, we started from the same Mature equilibrium at *E* = 1, held *E* at *E*_pre_ for *T*_pre_, and then switched to *E*_late_. The late-time mode depended on *E*_pre_, *E*_late_, and *τ*_*F*_ (Fig. 5A). At *τ*_*F*_ = 12.5, pre-conditioning at *E*_pre_ = 0.2 led to the Oscillatory mode, whereas *E*_pre_ = 4 led to the Mature mode after return to *E* = 1 (Fig. 5B). Varying *E*_pre_ and *T*_pre_ showed that the Mature mode remained the late-time outcome across a broad range of histories, whereas sufficiently long exposure to low or high *E*_pre_ led to the Low mode (Fig. 5C). Once *A* falls below *A*_sw_, the system cannot return to a high-adhesion mode (Supplementary Text S2). With the stiffness-history protocol fixed, increasing *τ*_*F*_ shifted the late-time mode from Low to Oscillatory and then to Disassembled (Fig. 5D). Thus, prior ECM exposure altered the internal state, leading to history-dependent adhesion-mode selection under the same final conditions, reflecting mechanobiological memory.

**FIG. 5.**
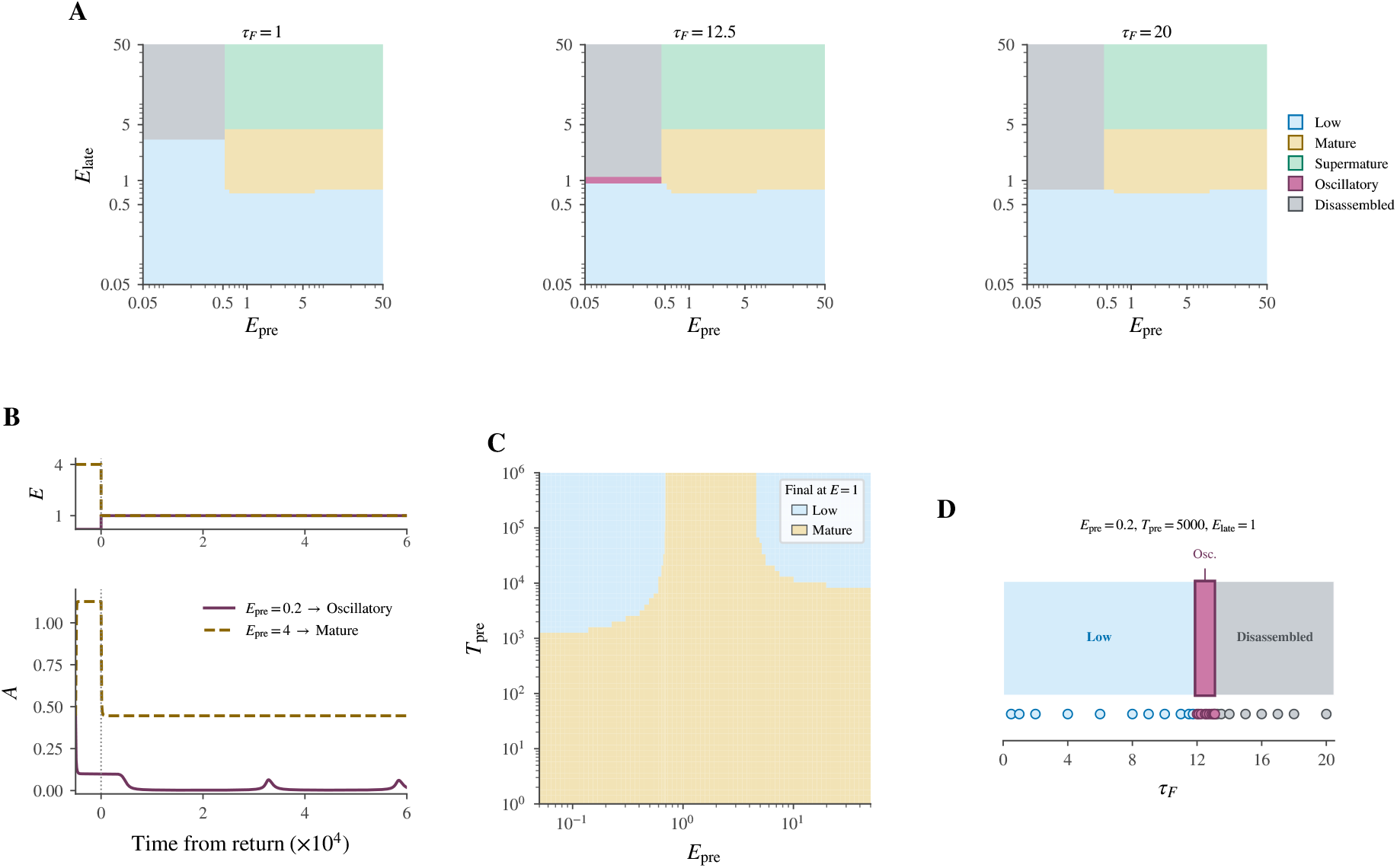
ECM history and force-response timescale shape adhesion-mode selection. All protocols start from the same Mature equilibrium at *E* = 1, with *τ*_*R*_ = *τ*_*S*_ = 1. (A) Late-time adhesion modes after preconditioning at *E*_pre_ for *T*_pre_ = 5000 and switching to *E*_late_, shown for *τ*_*F*_ = 1, 12.5, and 20. (B) Representative post-return responses at *E*_late_ = 1, *τ*_*F*_ = 12.5, and *T*_pre_ = 5000 for *E*_pre_ = 0.2 and 4. (C) Late-time adhesion mode after return to *E* = 1 as a function of *E*_pre_ and *T*_pre_, with *τ*_*F*_ = 1. (D) Dependence of mode selection on *τ*_*F*_ for *E*_pre_ = 0.2, *T*_pre_ = 5000, and *E*_late_ = 1.

## IV. DISCUSSION

To investigate how diverse adhesion modes emerge, we developed a mechanoadaptive model of focal adhesion– actomyosin coupling. The system supports Disassembled, Low, Mature, Supermature, and Oscillatory modes, with ECM stiffness organizing the stationary-mode structure and the force-response timescale reshaping dynamical stability (Fig. 6). Within multistable regimes, the internal state inherited from prior ECM exposure influences which adhesion mode is realized. These results indicate that a common mechanoadaptive system can give rise to state-dependent mechanical homeostasis, multi-stability, and mechanobiological memory, providing a framework for understanding diverse adhesion behaviors.

**FIG. 6.**
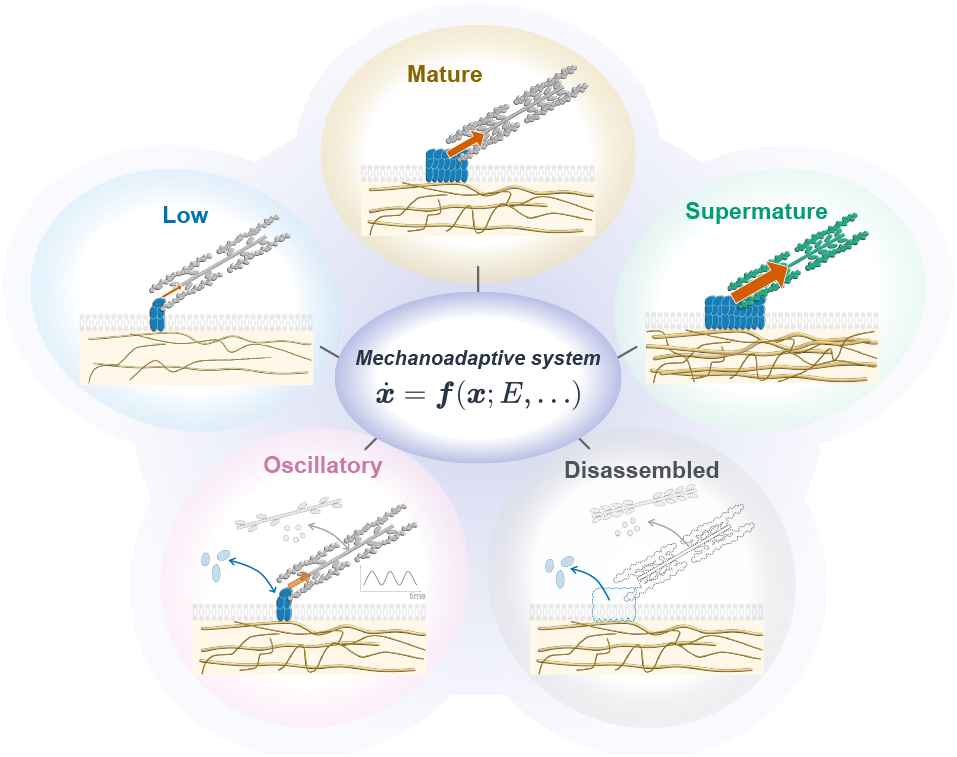
Diverse adhesion modes emerge from a common mechanoadaptive system. The schematic illustrates five adhesion modes of the focal adhesion–actomyosin system, with ECM stiffness shaping mode selection.

The adhesion modes broadly correspond to experimentally described focal-adhesion phenotypes in their structural and mechanical features. The Disassembled mode captures the loss of a force-bearing adhesion through disassembly [23]. Nascent adhesions and focal complexes share features with the Low mode [14, 15, 53, 54]. The Mature mode captures key features of force-bearing focal adhesions whose maturation is coupled to actomyosin-generated tension and stress-fiber organization [4–6, 55]. Large, high-stress adhesions in myofibroblasts are consistent with the Supermature mode, with their growth accompanying enhanced contractility and *α*-smooth muscle actin recruitment to stress fibers [19–21]. The Oscillatory mode represents a persistent adhesion with periodic mechanical activity, consistent with experimentally observed force oscillations and mechanochemical waves at adhesion sites [24–26].

Our results show that different ECM histories can lead to distinct adhesion modes under the same final stiffness. Previous studies have found that ECM stiffness history can affect focal-adhesion organization, actomyosin state, and cellular traction even after cells are placed in the same mechanical environment [56–58]. Our model suggests a dynamical mechanism for mechanobiological memory, in which prior ECM exposure alters the internal state and influences subsequent mode selection. Experimental responses to stiffness history depend on conditioning duration [56, 59, 60], consistent with the *T*_pre_ dependence of mode selection in our model. These findings suggest that the duration of ECM exposure can influence whether the system returns to its previous mode or transitions to a different one, reflecting both robustness and adaptability.

Different adhesion and contractile states can sustain distinct stress levels, motivating a state-dependent stress reference for mechanical homeostasis (or tensional homeostasis) [19, 45, 46]. Focal adhesions and stress fibers can adapt to load through growth, reinforcement, and repair [4, 11, 61, 62], supported by molecular processes including force-dependent binding, conformational changes, load sharing, and turnover [63–67]. In our model, adhesion stress closely tracks the adaptive target *S* at distinct levels in the stable Mature and Supermature states. The target thus serves as a reference for homeostatic regulation within a given state, while changes in the target itself can contribute to transitions between distinct states.

The force-response timescale can reorganize adhesion dynamics without changing the stationary equilibria. Increasing *τ*_*F*_ slows the adjustment of transmitted force toward its state-dependent capacity. Biologically, *τ*_*F*_ represents an effective timescale for force development and transmission through the stress-fiber–adhesion system and may depend on actomyosin activity, cytoskeletal mechanics, and adhesion–cytoskeleton coupling [50, 55, 68– 70]. Across a finite range of *τ*_*F*_, the Oscillatory mode coexists with the Mature equilibrium, expanding the dynamical repertoire at fixed ECM stiffness.

The coarse-grained model uses effective state variables and response timescales without resolving molecular kinetics. History-dependent adhesion-mode selection could be tested using reversibly tunable hydrogel substrates [71–74] to vary ECM preconditioning stiffness and duration, followed by time-resolved measurements of adhesion structure and traction [55, 75] and actomyosin dynamics [50] at a common final stiffness. Future extensions could incorporate spatial interactions among adhesions and slower adaptive processes, including transcriptional responses, long-term cytoskeletal reorganization, and persistent cell-state changes. The ECM could also be treated as a dynamical variable through matrix remodeling, including stiffening, maturation, and aging-associated changes. Overall, our results show that diverse adhesion modes can emerge within a common mechanoadaptive system combining homeostatic stress regulation and ECM-dependent force generation, with ECM stiffness history influencing mode selection.

## V. METHODS

### A. Stationary-state analysis and branch construction

We constructed stationary branches using an exact one-parameter reduction of the positive stationary-state conditions, with *A*^*\**^ as the branch coordinate. We write the system as

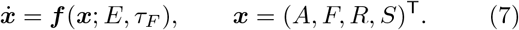

Stationary states satisfy ***f*** (***x***^*\**^; *E, τ*_*F*_ ) = **0**. For positive stationary states, the adhesion equation gives

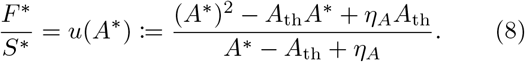

The remaining variables are reconstructed as *F* ^*\**^ = *S*^*\**^*u*(*A*^*\**^) and *R*^*\**^ = *χ*_*R*_*H*(*A*^*\**^; 1, *n*_*A→R*_)(*S*^*\**^ − 1), with either *S*^*\**^ = *S*_reg_(*A*^*\**^, *F* ^*\**^) or *S*^*\**^ = *S*_H_. The corresponding ECM-dependent force drive and stiffness are

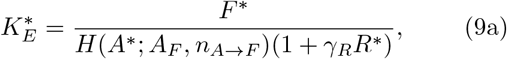

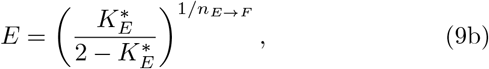

with 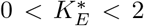. Only states with *A*^*\**^ *>* 0, *F* ^*\**^ *>* 0, *R*^*\**^ *>* 0, and 1 *< S*^*\**^ ≤ 1 + Δ_*S*_ were retained. For the reference parameter set and *E >* 0, this reduction exhaustively parameterizes the admissible positive stationary states.

Local stability was evaluated from the four-dimensional Jacobian ***J*** = *D*_***x***_***f*** and its spectral abscissa *α* = max_*j*_ Re *λ*_*j*_. Hyperbolic states with *α <* 0 were classified as locally asymptotically stable and those with *α >* 0 as unstable. The Disassembled state was treated separately because it is nonhyperbolic; nonlinear analysis established its local asymptotic stability within the physical state domain. Geometric folds of the stationary branches were located from d*E/*d*A*^*\**^ = 0. The exchange between the regulated and fixed-high *S* branches was determined separately from *S*_reg_ = *S*_H_. Spectral double-zero points were identified where two eigenvalues of the stationary-state Jacobian vanished simultaneously. Stress-target tracking of positive stationary states was quantified by the relative mismatch *ε*_*σ*_ = *F* ^*\**^*/*(*A*^*\**^*S*^*\**^) − 1 between the adhesion stress and the target. The stationary reduction, admissible state domain, Jacobian, and branch-classification procedure are given in Supplementary Text S2.

### B. Hopf bifurcation and periodic-orbit analysis

To characterize the emergence and stability of the Oscillatory mode, we analyzed Hopf bifurcations and periodic-orbit branches of the four-variable system. Hopf points were located when the Jacobian had a simple complex-conjugate pair

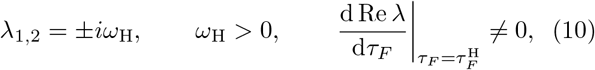

with no other eigenvalues on the imaginary axis. The transversality condition was evaluated with respect to *τ*_*F*_ at fixed *E*. Hopf criticality was assessed from the first Lyapunov coefficient *l*_1_ and the eigenvalue-crossing direction. With normalized phase *s* := *t/P*, a periodic orbit satisfies

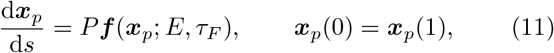

together with a phase condition. Periodic branches were continued in *τ*_*F*_ at fixed *E* and in *E* to map their organization in the (*E, τ*_*F*_ ) plane. Only periodic solutions satisfying *A >* 0, *F >* 0, *R >* 0, and 1 *< S <* 1 + Δ_*S*_ throughout the orbit were classified as positive-*R* Oscillatory modes.

Orbital stability was assessed using the variational equation

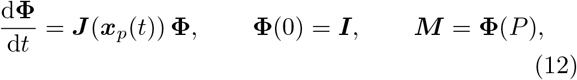

where **Φ**(*t*) is the fundamental matrix of the linearized dynamics along the periodic orbit and ***M*** is the corresponding monodromy matrix over one period *P* . The eigenvalues of ***M*** are the Floquet multipliers; after separating the neutral phase direction, a periodic solution was classified as orbitally stable when all nontrivial transverse multipliers satisfied |*µ*_*j*_| *<* 1. A limit point of cycles was assigned only when a periodic-branch turning point coincided with a nontrivial transverse multiplier crossing +1. Stationary–periodic coexistence was assigned only when a stationary equilibrium was locally stable and a periodic orbit was independently orbitally stable at the same parameter values. Further details are provided in Supplementary Text S3.

### C. Dynamical simulations and history-dependent mode selection

Representative dynamics and history-dependent mode selection were examined by direct integration of the four-variable system. All history protocols were initialized from the same Mature equilibrium at *E* = 1. At *t* = 0, ECM stiffness was switched to *E*_pre_ and subsequently to *E*_late_ at *t* = *T*_pre_, according to

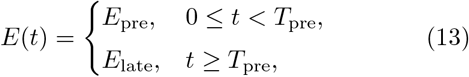

with continuous inheritance of the internal state, 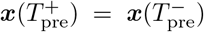. History-dependent mode selection was identified when distinct preconditioning protocols led to different late-time modes under the same *E*_late_, with all other model parameters held fixed.

Late-time stationary responses were assigned to the Disassembled, Low, Mature, or Supermature modes, whereas the Oscillatory mode required sustained oscillations in the positive-*R* domain. Damped oscillations and unresolved trajectories were not assigned to the Oscillatory class. These labels describe late-time response classes over the prescribed observation windows; attractor existence and stability were established independently. Further details of the history protocols and classification procedures are provided in Supplementary Text S4, with numerical implementation and acceptance criteria in Supplementary Text S5.

### D. Numerical implementation and verification

Computations used Python for nonlinear root finding, eigenvalue analysis, ODE integration, periodic boundary-value problems, and automatic differentiation to evaluate the first Lyapunov coefficient. Stationary states were verified using the vector-field residual and Jacobian spectrum. Periodic solutions were verified by periodic closure and the phase condition, with orbital stability assessed by Floquet analysis. Calculations near bifurcation or classification boundaries were repeated at tighter numerical tolerances. Software versions, solver settings, numerical tolerances, acceptance criteria, and reproducibility procedures are provided in Supplementary Text S5.

## DATA AND CODE AVAILABILITY

The numerical data and code used for the analyses, simulations, and figure generation will be made publicly available in a GitHub repository upon acceptance.

## AUTHOR CONTRIBUTIONS

E.M. and S.D. conceptualized the study. E.M. developed the mathematical model, performed the theoretical and computational analyses, and prepared the figures. S.D. supervised the project and contributed to the interpretation of the results. Both authors contributed to the original draft, reviewed and edited the manuscript, and acquired funding.

## DECLARATION OF GENERATIVE AI USE

During the preparation of this work, the authors used ChatGPT (OpenAI) to assist with language editing, literature exploration, code development and debugging, and mathematical and computational checks. All analyses, interpretations, and scientific conclusions were independently evaluated and verified by the authors, who take full responsibility for the content of the manuscript.

## SUPPLEMENTARY INFORMATION

### Supplementary Figures

**FIG. S1.**
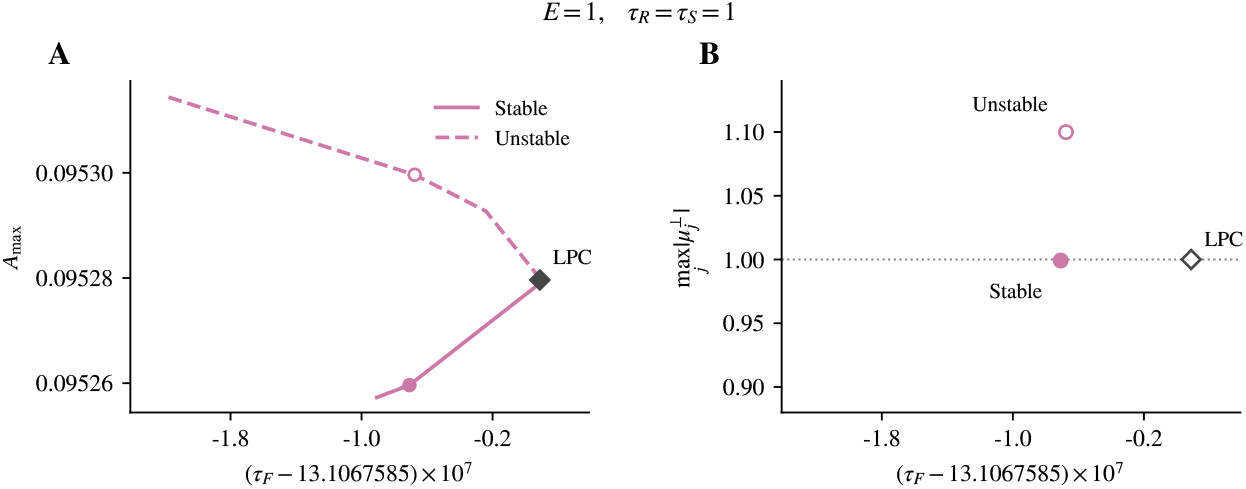
Local verification of the limit point of cycles at *E* = 1. (A) Local continuation of the positive-*R* periodic branch near the LPC, showing the stable and unstable branches meeting at the periodic-orbit fold. (B) Largest nontrivial transverse Floquet-multiplier modulus on the two sides of the same fold. The stable-side value lies below unity, whereas the unstable-side value exceeds unity, supporting a transverse +1 crossing at the LPC. The open LPC symbol in (B) marks the fold location as a reference and does not represent an independently evaluated Floquet multiplier exactly at the fold. Here, *τ*_*R*_ = *τ*_*S*_ = 1.

**FIG. S2.**
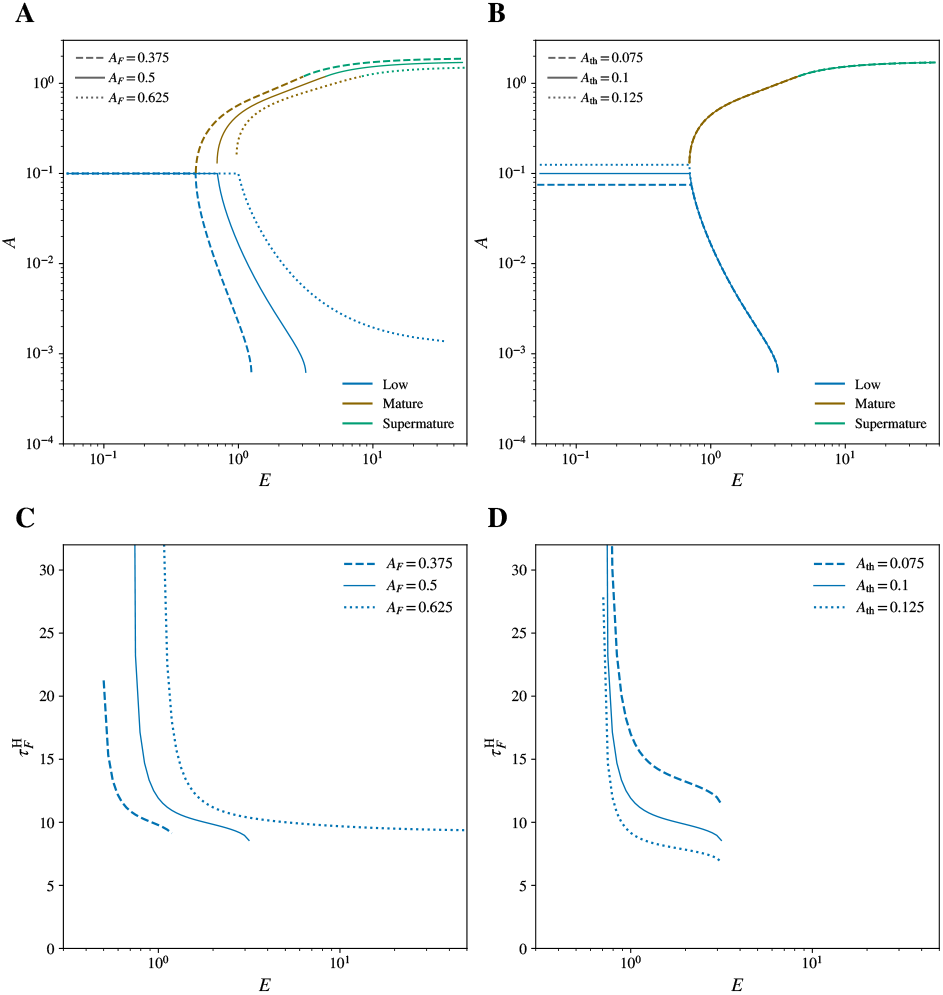
Robustness of stationary-state organization and Low-state Hopf instability under parameter variations. (A,B) Locally stable stationary branches for variations in *A*_*F*_ (A) and *A*_th_ (B), with *τ*_*F*_ = *τ*_*R*_ = *τ*_*S*_ = 1. Colors indicate the Low, Mature, and Supermature states, and line styles indicate the lower, reference, and upper parameter values. (C,D) Low-state Hopf loci for the corresponding variations in *A*_*F*_ (C) and *A*_th_ (D), with *τ*_*R*_ = *τ*_*S*_ = 1. The principal stationary-state organization and Low-state Hopf instability are preserved across the examined parameter variations, although their locations shift. All unvaried parameters were fixed at their reference values.

**FIG. S3.**
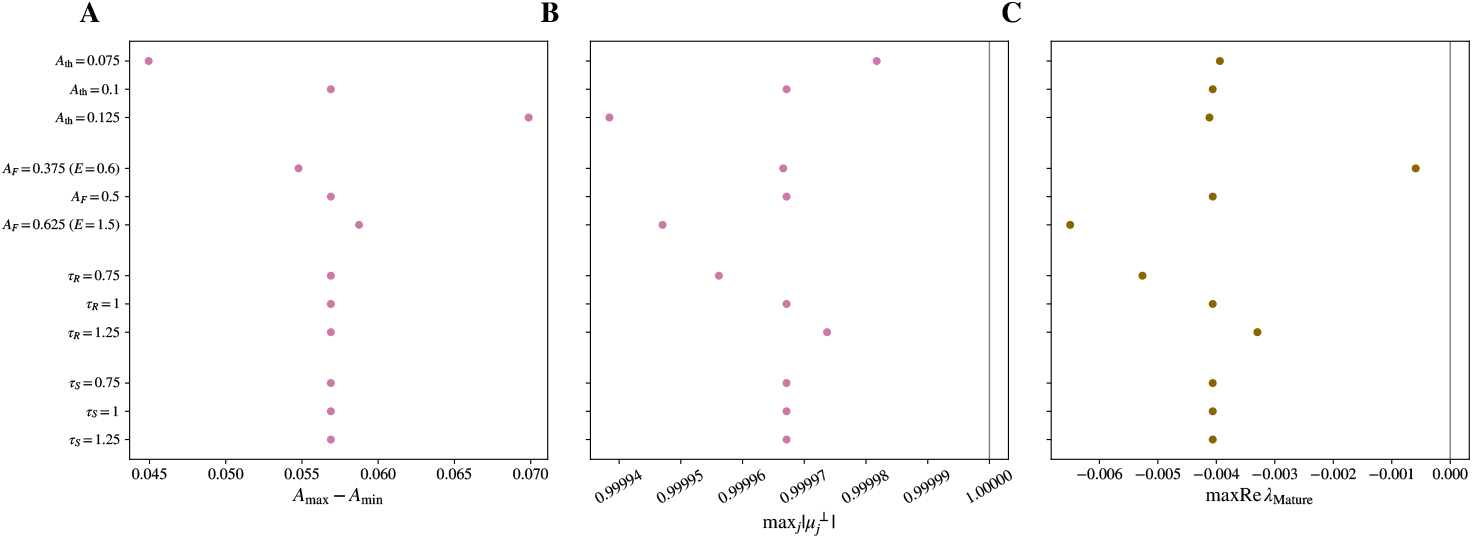
Persistence of stable oscillatory dynamics and Mature–Oscillatory coexistence under parameter variations. For variations in *A*_th_, *A*_*F*_, *τ*_*R*_, and *τ*_*S*_, each group shows the lower, reference, and upper parameter values. (A) Adhesion-amplitude range *A*_max_ *− A*_min_ of independently verified positive-*R* periodic solutions. (B) Largest nontrivial transverse Floquet-multiplier modulus, max_*j*_ 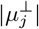, for the corresponding periodic solutions; values below unity indicate orbital stability. (C) Largest real part of the Jacobian eigenvalues, max Re *λ*_Mature_, of the independently verified Mature equilibrium at the same control point; values below zero indicate local stability. At the verified control points, stable periodic solutions coexist with stable Mature equilibria under all examined parameter variations. All points use *E* = 1 except *A*_*F*_ = 0.375 (*E* = 0.6) and *A*_*F*_ = 0.625 (*E* = 1.5), with the corresponding control points listed in Table S1. All unvaried parameters were fixed at their reference values.

### Supplementary Tables

**TABLE S1.** Verification of stable periodic solutions and Mature–Oscillatory coexistence under representative parameter variations. Each row reports a control point used for the periodic-orbit verification in Fig. S3. *P* is the period of the verified positive-*R* periodic solution, *A*_max_ *− A*_min_ is its adhesion-amplitude range, max_*j*_ 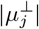 is the largest nontrivial transverse Floquet-multiplier modulus, and max Re *λ*_Mature_ is the spectral abscissa of the Mature equilibrium at the same parameter values. Orbital stability requires max_*j*_ 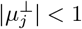, whereas local stability of the Mature equilibrium requires max Re *λ*_Mature_ *<* 0. The reference condition is listed once, although the same reference solution is displayed in each parameter group in Fig. S3. All unlisted parameters were fixed at their reference values. Full-precision numerical values are retained in the reproducibility package.

| Varied parameter | Value | $E$ | $\tau_F$ | $P$ | $A_{\max} - A_{\min}$ | $\max_j \vert \mu_j^\perp\vert $ | $\max \text{Re } \lambda_{\text{Mature}}$ |
| --- | --- | --- | --- | --- | --- | --- | --- |
| Reference | – | 1 | 12.5 | $2.42 \times 10^4$ | 0.0569 | 0.999967 | $-4.062 \times 10^{-3}$ |
| $A_{\text{th}}$ | 0.075 | 1 | 17.86 | $2.97 \times 10^4$ | 0.0449 | 0.999982 | $-3.939 \times 10^{-3}$ |
| $A_{\text{th}}$ | 0.125 | 1 | 9.624 | $2.21 \times 10^4$ | 0.0699 | 0.999938 | $-4.118 \times 10^{-3}$ |
| $A_F$ | 0.375 | 0.6 | 12.74 | $1.96 \times 10^4$ | 0.0548 | 0.999967 | $-5.902 \times 10^{-4}$ |
| $A_F$ | 0.625 | 1.5 | 13.39 | $1.61 \times 10^4$ | 0.0587 | 0.999947 | $-6.499 \times 10^{-3}$ |
| $\tau_R$ | 0.75 | 1 | 12.5 | $2.42 \times 10^4$ | 0.0569 | 0.999956 | $-5.264 \times 10^{-3}$ |
| $\tau_R$ | 1.25 | 1 | 12.5 | $2.42 \times 10^4$ | 0.0569 | 0.999974 | $-3.298 \times 10^{-3}$ |
| $\tau_S$ | 0.75 | 1 | 12.5 | $2.42 \times 10^4$ | 0.0569 | 0.999967 | $-4.062 \times 10^{-3}$ |
| $\tau_S$ | 1.25 | 1 | 12.5 | $2.42 \times 10^4$ | 0.0569 | 0.999967 | $-4.061 \times 10^{-3}$ |

**TABLE S2.**
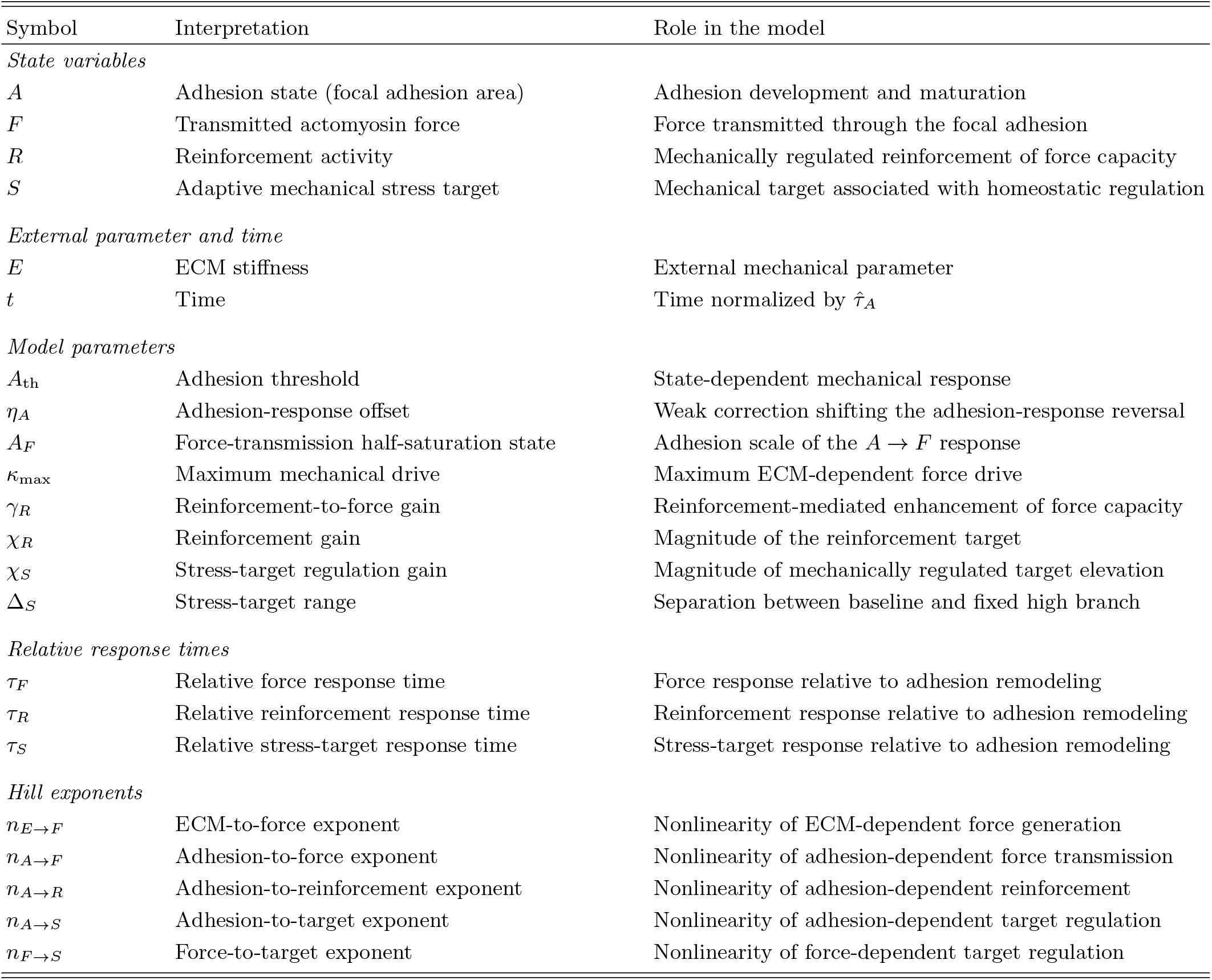
State variables and parameters of the dimensionless four-variable model.

## SUPPLEMENTARY TEXT S1. DIMENSIONAL FORMULATION AND NONDIMENSIONALIZATION

The model is formulated in terms of four coarse-grained variables describing a coupled focal adhesion–actomyosin system. Because the model analyzed in the main text is nondimensional, we first introduce a dimensional formulation and then derive the nondimensional system.

### Dimensional formulation

We denote dimensional quantities by hats in this section. The variables *Â*, 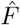, *R*, and *Ŝ* represent focal adhesion area, transmitted actomyosin force, reinforcement activity, and the adaptive mechanical stress target, respectively; *R* is dimensionless. The ratio 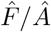 represents the average mechanical stress supported by the focal adhesion and is proportional to the average force per force-bearing adhesion molecule when their areal density is approximately constant.

Saturating regulatory responses are represented by the Hill function

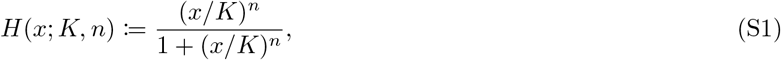

where *K* denotes the half-saturation value and *n* controls the response nonlinearity.

We construct the adhesion-remodeling law in multiplicative form so that remodeling vanishes continuously as the adhesion disappears. Mechanical force promotes focal adhesion growth, while adhesion maturation is accompanied by changes in Rho-family GTPase signaling and actomyosin contractility [1–5]. We represent the corresponding maturation-dependent change in mechanical feedback through the threshold *Â*_th_. Force and area are approximately coupled for individual focal adhesions, whereas distinct adhesion states can support different characteristic stress levels [6–8]. These observations motivate the force–adhesion mismatch 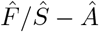, giving the leading remodeling structure

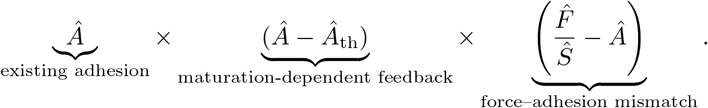

Because this term vanishes identically at *Â* = *Â*_th_, we add the weak perturbation 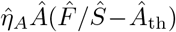 to remove this exact pinning. Introducing a reference adhesion area *Â*_0_ and response-time scale 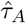, the dimensional adhesion dynamics are

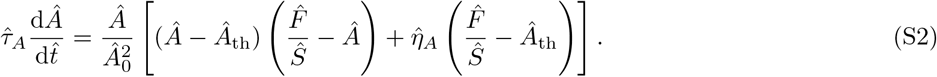

Here, 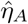 has dimensions of area and is assumed to satisfy 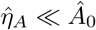.

Focal adhesion development is coupled to actomyosin contractility [9, 10], cellular force generation depends on ECM stiffness [11, 12], and reinforcement of the contractile machinery can further increase force generation [8, 13].

We therefore define the ECM-dependent mechanical drive as

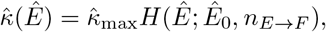

and the effective force-generating capacity as

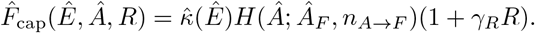

Here, *Ê*_0_ and *Â*_*F*_ are the corresponding half-saturation scales, *n*_*E→F*_ and *n*_*A→F*_ are Hill exponents, 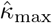 sets the maximum drive, and *γ*_*R*_ is the reinforcement-to-force gain. The dimensional force dynamics are

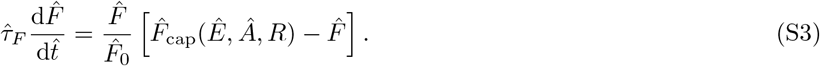

Here, 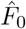 is the reference force scale defined below.

Mechanically reinforced stress fibers involve tension- and adhesion-dependent processes that enhance contractility, including recruitment of *α*-smooth muscle actin and phosphorylation of myosin regulatory light chain [8, 14, 15]. We represent reinforcement as an adhesion-dependent response to elevation of the adaptive stress target above baseline,

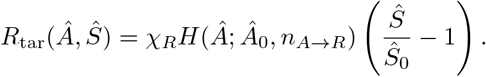

The reinforcement dynamics are written in multiplicative form to preserve the unreinforced state *R* = 0,

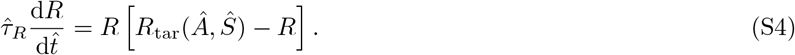

Here, *χ*_*R*_ is the reinforcement gain and *n*_*A→R*_ controls its adhesion dependence.

Distinct focal adhesion states can support different characteristic stress levels, and cellular mechanical homeostasis involves regulation around characteristic mechanical operating levels [8, 16, 17]. Multiple mechanical set points have also been proposed theoretically [18]. We therefore introduce a fixed high branch *Ŝ*_H_ and a mechanically regulated branch *Ŝ*_reg_ (*Â*, 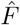),

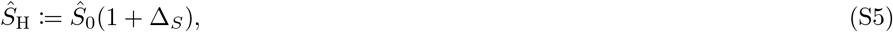

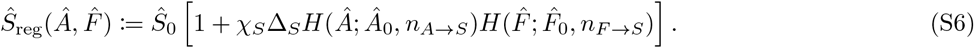

Here, Δ_*S*_ sets the separation between the baseline and fixed-high branches, *χ*_*S*_ controls the regulated target elevation, and *n*_*A→S*_ and *n*_*F →S*_ are the corresponding Hill exponents. The stress-target dynamics are

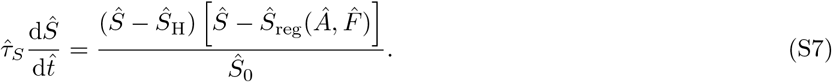

Both *Ŝ*_H_ and *Ŝ*_reg_ are stationary branches. The physical interval *Ŝ*_0_ ≤ *Ŝ* ≤ *Ŝ*_H_ is forward invariant and ensures a non-negative reinforcement target. The positive parameters 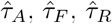 and 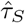 set the corresponding kinetic scales.

### Nondimensionalization

We choose *Â*_0_, *Ŝ*_0_, *Ê*_0_, and 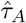 as the reference scales for adhesion area, stress, ECM stiffness, and time, respectively. The corresponding force scale is 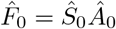, so that 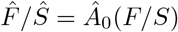. We define the dimensionless variables and time by

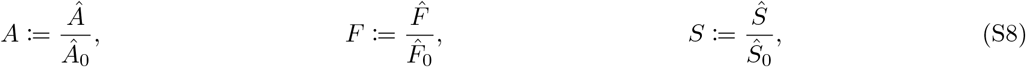

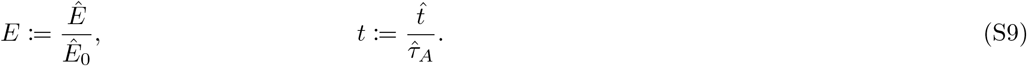

The reinforcement state *R* is already dimensionless.

The remaining dimensional parameters give

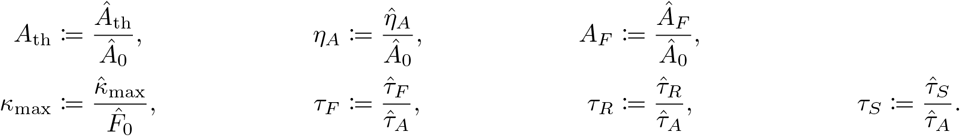

The parameters *γ*_*R*_, *χ*_*R*_, *χ*_*S*_, and Δ_*S*_, together with all Hill exponents, are already dimensionless.

### Dimensionless model and notation

Substitution of the above definitions gives

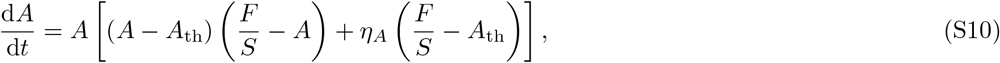

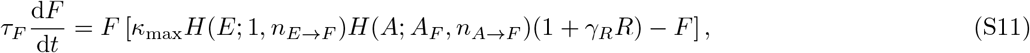

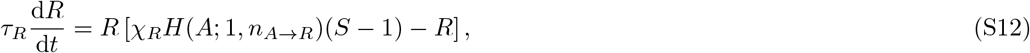

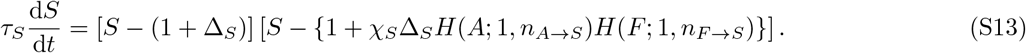

For the reference model, we set *κ*_max_ = 2 and define

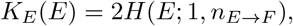

so that

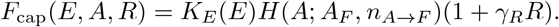

The common Hill function *H* is given by Eq. (S1). The dimensionless state vector is ***x***(*t*) = (*A*(*t*), *F* (*t*), *R*(*t*), *S*(*t*))^T^, with physical domain *A* ≥ 0, *F* ≥ 0, *R* ≥ 0, and 1 ≤ *S* ≤ 1 + Δ_*S*_; *E* ≥ 0 is treated as an external mechanical parameter. Unless otherwise stated, all response-time, gain, range, half-saturation, and Hill-exponent parameters are positive. The dimensionless variables and parameters are summarized in Table S2.

## SUPPLEMENTARY TEXT S2. STATIONARY STATES AND LOCAL STABILITY

### Stationary states and physical state domain

We consider the dimensionless system defined in Supplementary Text S1 on the physical domain

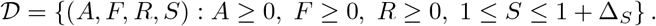

The planes *A* = 0, *F* = 0, and *R* = 0 are invariant. At *S* = 1,

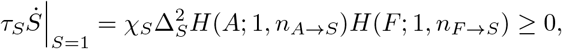

whereas *S* = 1 + Δ_*S*_ is invariant; hence *D* is forward invariant.

The boundary state *D* = (0, 0, 0, 1) is an exact stationary solution for all *E* ≥ 0 and is denoted Disassembled. For *E >* 0, equilibria with *A >* 0 and *F* = 0 are transversely unstable because the linearized force growth rate is *F*_cap_*/τ*_*F*_ *>* 0. Equilibria with *A >* 0, *F >* 0, and *R* = 0 satisfy *S >* 1 and are transversely unstable because the linearized reinforcement growth rate is *R*_tar_*/τ*_*R*_ *>* 0. The remaining equilibrium on the extinction boundary, (*A, F, R, S*) = (0, 0, 0, *S*_H_), is unstable in the *S* direction because 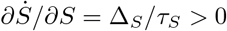. Thus, for *E >* 0, all equilibria on an extinction face *AFR* = 0 are unstable except *D*. Positive equilibria on the invariant boundary *S* = *S*_H_ are treated separately below. We therefore restrict the positive stationary-state analysis to *A >* 0, *F >* 0, and *R >* 0. The superscript * is omitted for stationary quantities throughout this Supplementary Text.

For a positive stationary state, the adhesion equation gives

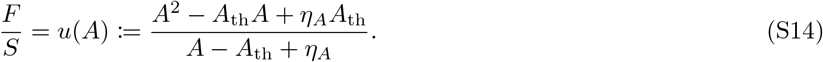

The denominator vanishes at *A*_sw_ = *A*_th_ − *η*_*A*_. For 0 *< η*_*A*_ *< A*_th_*/*4, the numerator has the positive roots

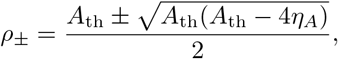

and *F/S >* 0 restricts positive stationary states to

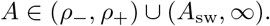

The pole at *A* = *A*_sw_ separates the low- and high-adhesion components of the reduced stationary representation, although the original dynamical system is regular there. At this switching state,

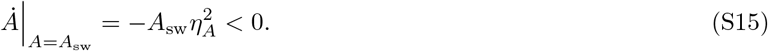

Hence *A* = *A*_sw_ can be crossed only toward decreasing *A*, and *A* ≤ *A*_sw_ is forward invariant. For each admissible *A*,

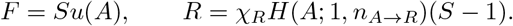

The *S* equation gives either the regulated (REG) family,

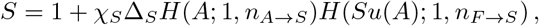

or the fixed-high (SAT) family,

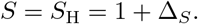

For the reference values *n*_*F →S*_ = 2, *n*_*A→S*_ = 2, *χ*_*S*_ = 2, and Δ_*S*_ = 1, define

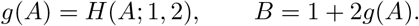

The REG condition reduces to

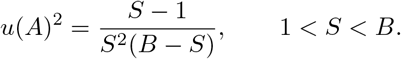

Because the right-hand side is strictly increasing for 1 *< S < B*, there is at most one positive REG root for each admissible *A*; only roots satisfying 1 *< S* ≤ 1 + Δ_*S*_ were retained.

The force equation determines the ECM-dependent drive and stiffness through

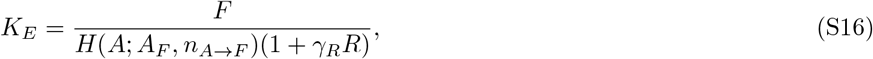

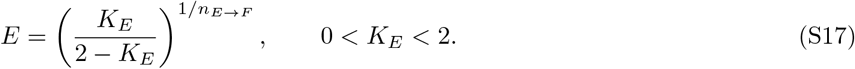

Equations (S14) and (S17) therefore parameterize the positive stationary branches by *A*. For the reference parameter set and *E >* 0, every admissible positive equilibrium belongs to either the REG or SAT family and is reconstructed uniquely subject to the physical-domain and 0 *< K*_*E*_ *<* 2 constraints.

### Linear stability

Local stability was evaluated from the full four-dimensional Jacobian

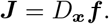

For compactness, define

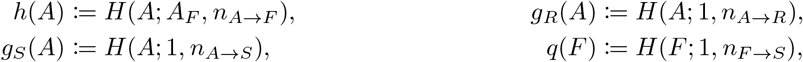

and let primes denote derivatives with respect to the function argument. With *u* = *F/S* and *d* = *A* − *A*_th_ + *η*_*A*_, the nonzero Jacobian entries are

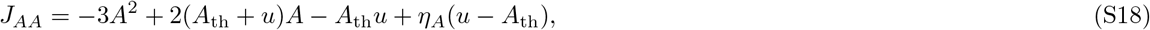

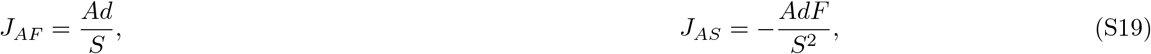

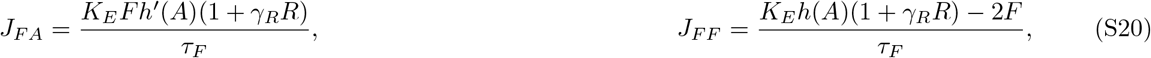

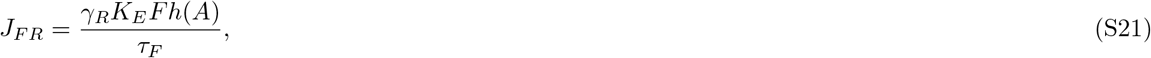

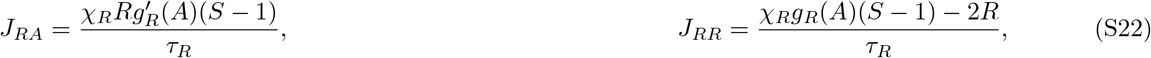

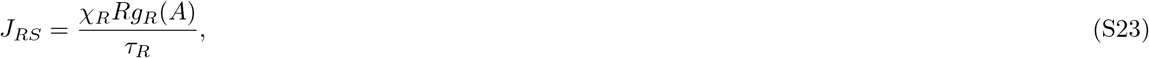

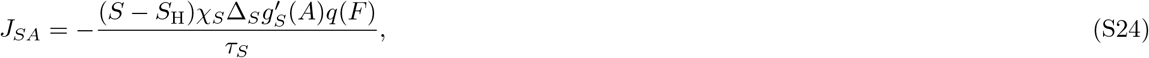

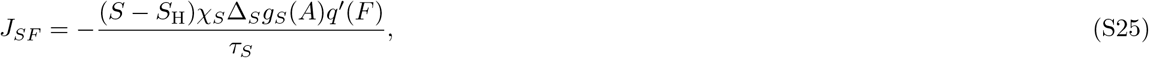

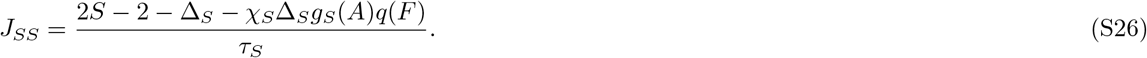

The remaining entries *J*_*AR*_, *J*_*F S*_, *J*_*RF*_, and *J*_*SR*_ vanish.

Local stability was classified using the spectral abscissa

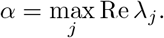

Hyperbolic equilibria with *α <* 0 were classified as locally asymptotically stable and those with *α >* 0 as unstable. Nonhyperbolic states were treated separately. The response-time parameters do not affect stationary-state geometry but can alter local stability; the classification in Fig. 2 uses *τ*_*F*_ = *τ*_*R*_ = *τ*_*S*_ = 1. Numerical procedures and treatment of eigenvalues near the imaginary axis are given in Supplementary Text S5.

### Local stability of the Disassembled state

The Disassembled state is nonhyperbolic. At

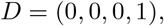

the Jacobian spectrum is

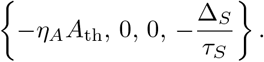

Thus the *A* and *S* directions are linearly stable, whereas the *F* and *R* directions require nonlinear analysis.

Let

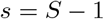

and consider

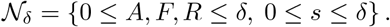

Because the factor multiplying *A* equals −*η*_*A*_*A*_th_ *<* 0 at *D*, sufficiently small *δ >* 0 gives

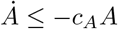

for some *c*_*A*_ *>* 0 throughout *N*_*δ*_. For the reference exponent *n*_*A→F*_ = 1.2 *>* 1,

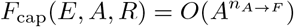

for bounded *R* and fixed finite *E*. Similarly,

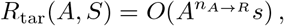

and

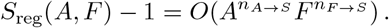

Because *n*_*A→F*_ *>* 1 and the remaining Hill responses vanish at least quadratically near the origin, *δ* can be chosen sufficiently small so that the vector field points inward on the upper *F, R*, and *s* boundaries of N_*δ*_, while the lower boundaries are invariant or inward-pointing. Hence *N*_*δ*_ is forward invariant, and

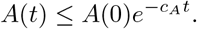

Define

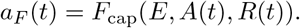

Since *A*(*t*) decays exponentially and *R*(*t*) remains bounded, *a*_*F*_ (*t*) is nonnegative and integrable. For *F* (0) *>* 0, the reciprocal variable *y*_*F*_ = 1*/F* satisfies

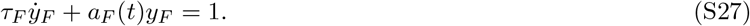

Hence

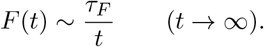

If *F* (0) = 0, invariance of *F* = 0 gives *F* (*t*) ≡ 0.

Similarly, let

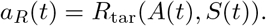

The function *a*_*R*_(*t*) is nonnegative and integrable, and for *R*(0) *>* 0, *y*_*R*_ = 1*/R* satisfies

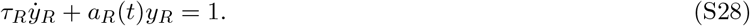

Therefore

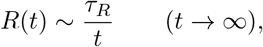

whereas *R*(0) = 0 implies *R*(*t*) ≡ 0.

Finally, define

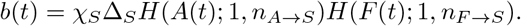

The stress-target dynamics become

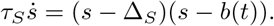

Because *A*(*t*) decays exponentially, *b*(*t*) also decays exponentially. For sufficiently small *δ <* Δ_*S*_, comparison with the corresponding stable linear equation gives

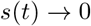

exponentially.

Thus,

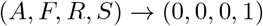

for all sufficiently small perturbations of *D* within the physical domain. The Disassembled state is therefore locally asymptotically stable relative to the physical domain. For positive initial *F* and *R*, the nonhyperbolic directions decay algebraically as *F* (*t*) ∼ *τ*_*F*_ */t* and *R*(*t*) ∼ *τ*_*R*_*/t*, whereas *A* and *S* − 1 decay exponentially.

### Stationary landmarks and adhesion-mode classification

A geometric fold of a REG or SAT branch was defined by

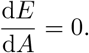

The REG–SAT exchange instead occurs when

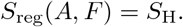

At the exchange, the two stationary *S* roots coincide and the *S* row of the Jacobian vanishes, producing a zero eigenvalue. Thus the REG–SAT exchange is nonhyperbolic but distinct from a geometric fold. Geometric folds were identified from the reduced stationary branches and checked against the original four-variable equations and Jacobian spectrum.

The stable low-area REG, high-area REG, and high-area SAT branches are denoted Low, Mature, and Supermature, respectively; the exact boundary state *D* = (0, 0, 0, 1) is denoted Disassembled. Unstable positive branches are retained in the stationary diagrams but are not classified as stable adhesion modes.

At a positive stationary state, the relative mismatch between the adhesion stress and the adaptive target is

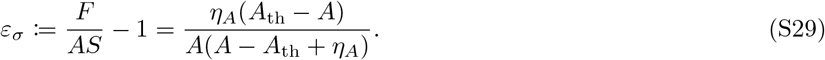

For the stable stationary branches at the reference response timescales, the maximum absolute mismatch is 0.0773% for the Mature mode and 0.00838% for the Supermature mode. These high-adhesion modes therefore closely track their mechanical stress target. The Low branch does not exhibit uniformly close target tracking and is not included in this homeostatic interpretation.

## SUPPLEMENTARY TEXT S3. HOPF BIFURCATIONS AND PERIODIC-ORBIT ANALYSIS

### Hopf bifurcation and criticality

Let ***x***^*\**^ be a stationary solution of the full four-variable system with Jacobian ***J*** ^*\**^ = *D*_***x***_***f*** (***x***^*\**^; *E, τ*_*F*_ ). At fixed ECM stiffness *E*, a Hopf point was identified when ***J*** ^*\**^ possessed a simple complex-conjugate eigenvalue pair satisfying

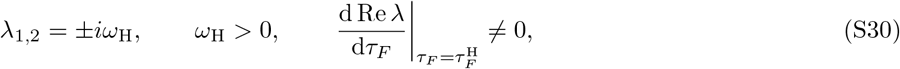

with no other eigenvalues on the imaginary axis.

Hopf criticality was determined from the first Lyapunov coefficient. Let ***q*** and ***p*** denote the right and adjoint critical eigenvectors satisfying

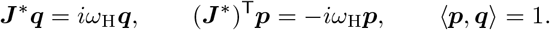

Expanding the vector field as

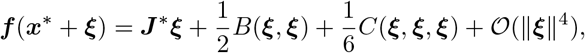

where *B* and *C* are the quadratic and cubic multilinear forms generated by the second and third derivatives of the vector field, we evaluated

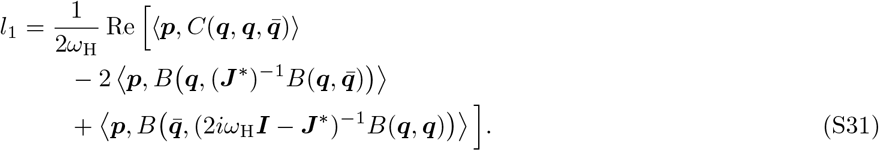

Under this convention, *l*_1_ *<* 0, together with the eigenvalue-crossing direction, identifies a supercritical Hopf bifurcation. Numerical differentiation and scaling checks are described in Supplementary Text S5.

The spectral double-zero point in the stationary-state stability map was treated only as a point at which two eigenvalues simultaneously vanish; no nonlinear codimension-two classification was assigned.

### Periodic-orbit continuation and Floquet stability

Periodic solutions of period *P* were computed from the full four-variable system using the normalized phase *s* = *t/P*,

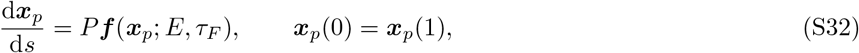

together with a phase condition. At fixed *E*, periodic solutions were continued in *τ*_*F*_, and continuation across *E* was used to track the branch in the (*E, τ*_*F*_ ) plane. Positive-*R* Oscillatory modes were restricted to cycles satisfying

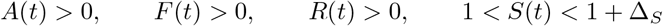

throughout the period.

For an infinitesimal perturbation *δ****x*** about a nonconstant periodic solution,

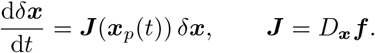

Defining *δ****x***(*t*) = **Φ**(*t*)*δ****x***(0) gives the fundamental-matrix equation

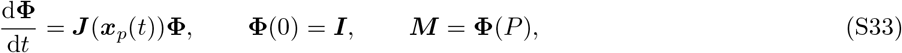

or componentwise,

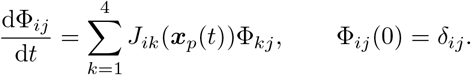

The eigenvalues *µ*_*j*_ of the monodromy matrix ***M*** are the Floquet multipliers.

Because the system is autonomous, 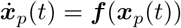 satisfies the variational equation, and

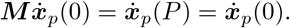

Thus every nonconstant periodic orbit has a neutral phase multiplier *µ*_ph_ = 1 in exact arithmetic. The numerical boundary-value and variational calculations used scaled logarithmic coordinates

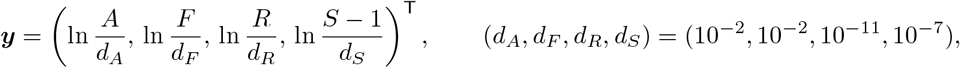

with

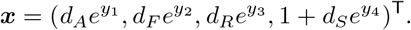

Writing 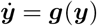, the infinitesimal coordinate transformation is

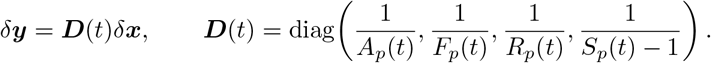

Let **Ψ**(*t*) denote the logarithmic-coordinate fundamental matrix and **Ψ**_n_(*s*) = **Ψ**(*Ps*). The numerically integrated variational system was

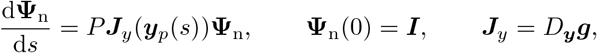

with

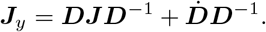

The physical- and logarithmic-coordinate fundamental matrices satisfy

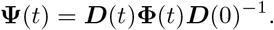

Because ***D***(*P* ) = ***D***(0),

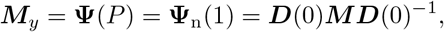

so ***M***_*y*_ and ***M*** have identical Floquet multipliers.

The three transverse multipliers were computed separately from the full monodromy spectrum. After choosing the origin at a sampled phase with maximum logarithmic tangent speed, define

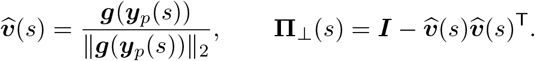

Let ***Q***_0_ be an orthonormal 4 × 3 basis for the null space of 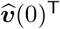. For *s*_*i*_ = *i/N* and Δ*s* = 1*/N*, the implemented propagation was

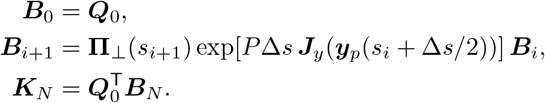

Projection was applied after each short-step matrix exponential without renormalization or reorthogonalization. The eigenvalues of ***K***_*N*_ approximate the three nontrivial Floquet multipliers as the short-step approximation is refined. This construction removes the tangent direction geometrically rather than by deleting an eigenvalue near +1.

The phase separation was checked by step refinement and the logarithmic tangent-invariance defect

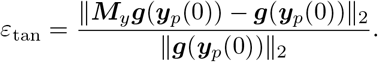

A periodic orbit was classified as orbitally stable when

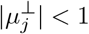

for all three transverse multipliers.

An LPC was identified from a local turning point of *τ*_*F*_ along the amplitude-parameterized periodic branch together with positive real nontrivial transverse multipliers bracketing +1 on its stable and unstable sides. Numerical tolerances and verification procedures are given in Supplementary Text S5.

### Periodic-orbit topology and coexistence

At *E* = 1, the Low equilibrium undergoes a Hopf bifurcation at

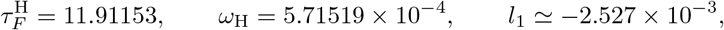

which is supercritical. The positive-*R* periodic branch is orbitally stable over a finite interval of *τ*_*F*_, up to an LPC near 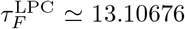. The LPC is supported by the periodic-branch turning point and the transverse +1 crossing (Fig. S1).

At *τ*_*F*_ = 12.5, the representative positive-*R* periodic orbit has

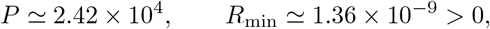

and largest transverse Floquet-multiplier modulus approximately 0.999967, confirming orbital stability. This orbit coexists with the locally stable Mature equilibrium at the same parameter values (Fig. 3C and Fig. 4G).

Across *E*, the verified positive-*R* periodic branch defines the Oscillatory region shown in Fig. 3A. The higher-*E* outer boundary is shown as an LPC only where both branch turning and a transverse +1 crossing were verified. On the lower-*E* side, periods increased strongly near the numerical limit of continuation; the termination mechanism was therefore left unresolved rather than assigned a specific global bifurcation.

## SUPPLEMENTARY TEXT S4. HISTORY-DEPENDENT MODE SELECTION

### History protocol and mode classification

History-dependent responses were computed by direct integration of the full four-variable model under prescribed changes in ECM stiffness, with the internal state inherited continuously across each switch. Late-time responses were assigned to the Disassembled, Low, Mature, Supermature, Oscillatory, or unresolved classes using the operational criteria specified in Supplementary Text S5. Stationary modes are defined in Supplementary Text S2, and representative Oscillatory responses were cross-checked against the positive-*R*, orbitally stable periodic solutions described in Supplementary Text S3.

All protocols were initiated from the validated Mature stationary state at *E* = 1,

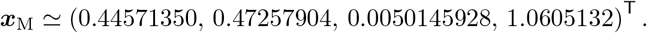

Preconditioning was imposed as

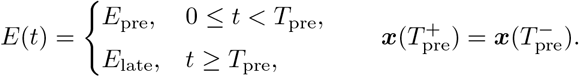

Unless otherwise stated, *τ*_*R*_ = *τ*_*S*_ = 1. History-dependent selection was identified when different preconditioning protocols produced different late-time modes under the same *E*_late_ and otherwise identical model parameters.

### Figure 5 protocols

For Fig. 5A, *T*_pre_ = 5000, and both *E*_pre_ and *E*_late_ were sampled on the same 35-point grid from 0.05 to 50. The grid was obtained by inserting geometric midpoints between adjacent values of

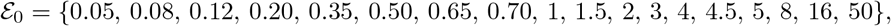

giving 35 × 35 = 1225 histories for each of *τ*_*F*_ = 1, 12.5, and 20. For Fig. 5B,

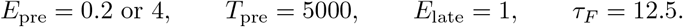

The final mode was evaluated over 3.6 × 10^5^ time units after return to *E* = 1; the plotted traces show the first 6 × 10^4^ time units.

For Fig. 5C, *E*_pre_ and *T*_pre_ were varied before return to *E*_late_ = 1, with *τ*_*F*_ = *τ*_*R*_ = *τ*_*S*_ = 1. The *E*_pre_ grid contained 72 logarithmically spaced values from 0.05 to 50 supplemented by

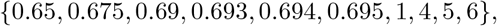

giving 82 unique values. The *T*_pre_ grid contained 60 logarithmically spaced values from 1 to 10^6^ together with *T*_pre_ = 5000, giving 61 values and 5002 histories in total. The resulting (*E*_pre_, *T*_pre_) map reports the late-time mode selected after return to *E* = 1; it does not represent a post-return memory lifetime.

For Fig. 5D,

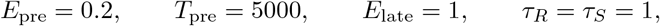

while *τ*_*F*_ was sampled at

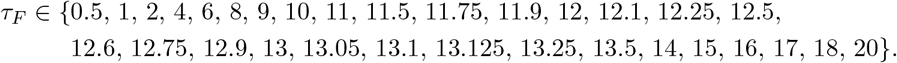

Sampling was denser near transitions involving the Oscillatory mode. The response at *τ*_*F*_ = 11.9 remained a slowly decaying nonstationary transient over the extended observation interval and was retained as unresolved. The colored intervals in Fig. 5D therefore summarize classifications at the sampled *τ*_*F*_ values and are not interpreted as bifurcation boundaries.

## SUPPLEMENTARY TEXT S5. NUMERICAL IMPLEMENTATION AND VERIFICATION

### Stationary states, bifurcations, and periodic solutions

Numerical calculations were performed in Python 3.12.14 with NumPy 2.3.5, SciPy 1.17.0, pandas 2.2.3, Matplotlib 3.10.8, and CasADi 3.7.2. Unless otherwise stated, all calculations used the four-variable model and the reference structural parameter set defined in the main text.

Stationary solutions were constructed from the reduced equations in Supplementary Text S2. Scalar equations were solved by bracketed Brent root finding with xtol = 2 × 10^*−*15^ and rtol = 10^*−*14^; geometric folds were refined by bounded one-dimensional minimization. Each reconstructed state was substituted into the original vector field. The final Figure 2 data had a maximum infinity-norm residual below 5 × 10^*−*12^, with refined values reaching approximately 5 × 10^*−*15^.

Local stability was determined from the full Jacobian spectrum. Cases with spectral abscissa magnitude below 10^*−*9^ were re-evaluated at 60- and 90-digit precision using *w* = *S* − 1 and diagonal similarity scaling. A stability sign was accepted only when it was consistent across precision levels and separated from numerical error. Geometric folds, the REG–SAT exchange, Hopf points, and the spectral double-zero point were evaluated from their respective defining conditions. The Disassembled state was treated analytically as described in Supplementary Text S2.

Hopf points were located by tracking and refining the zero crossing of the real part of the critical complex-conjugate eigenvalue pair. At *E* = 1, the Hopf value of *τ*_*F*_ was refined with absolute root tolerance 10^*−*13^. The first Lyapunov coefficient was evaluated using CasADi automatic differentiation of a diagonally scaled vector field. The representative scaling was

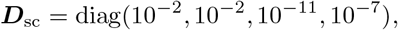

with the fourth coordinate corresponding to *S* − 1. Alternative diagonal scalings gave the same sign of *l*_1_.

Periodic solutions were computed by collocation using the formulation in Supplementary Text S3. Positive periodic solutions were represented in logarithmic coordinates for *A, F, R*, and *S* − 1. The waveform, period, and, where required, *τ*_*F*_ were solved together with a phase condition and continued from previously converged solutions. The positive-*R* periodic orbit at *E* = 1 and *τ*_*F*_ = 12.5 was refined with boundary-value tolerance 3 × 10^*−*9^.

Orbital stability was evaluated from the logarithmic-coordinate variational system using the implicit Radau method with

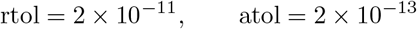

for the reference orbit. The full monodromy phase multiplier differed from unity by approximately 8.2 × 10^*−*12^, the largest transverse multiplier was 0.999967, and the tangent-invariance error was 3.3 × 10^*−*12^. Candidate LPCs were recalculated at tighter numerical settings and retained only when a periodic-branch turning point was accompanied by a transverse Floquet multiplier crossing +1.

Representative trajectories in Fig. 4 were integrated with SciPy solve ivp using BDF with

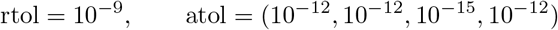

for (*A, F, R, S*) and max step = *t*_end_*/*900. Displayed trajectories were sampled on an evenly spaced output grid. Oscillatory panels used the independently verified positive-*R* periodic solution.

For the gray streamlines in Fig. 4F,G, the conditional 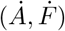 field was evaluated at fixed *R* and *S*. Panel F used the component-wise midpoint of the Low and Mature equilibria, whereas panel G used the midpoint of the Mature equilibrium and the arithmetic mean of the stored 1400-sample periodic orbit.

### ECM-history calculations

The history calculations in Fig. 5 used direct integration of the full model. For panels A and C,

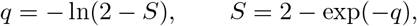

was used for 1 ≤ *S <* 2 to avoid numerical rounding onto the invariant *S* = 2 boundary. The transformed state was inherited continuously across ECM switches, and reported states were converted back to (*A, F, R, S*).

For panels A, B, and D, the physical-state residual was

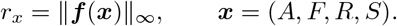

For panel C, the transformed residual was

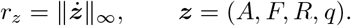

For panel A and the protocol-check and additional points in panel D, endpoint drift was

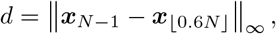

where *N* denotes the number of concatenated stored post-switch samples. For panel B and the panel-D points evaluated using the same procedure as panel B, the maximum component-wise late-time amplitude was

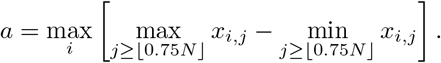

Component-wise division is used for the scaled vector quantities below. For panel C, the scaled drift over an integration stage [*t*_*a*_, *t*_*b*_] was

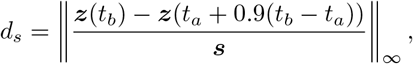

with

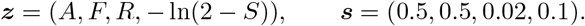

Oscillation persistence was quantified using

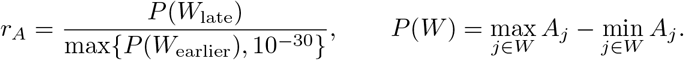

For panel A and the protocol-check and additional points in panel D,

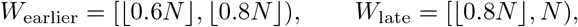

whereas panel B and the panel-D points evaluated using the same procedure as panel B used

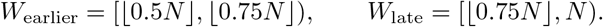

Duplicate stage-start samples were omitted when concatenating staged integrations. The focused panel-A rechecks used the corresponding 60–80% and final-20% sample windows within the current 701-point integration stage. Peaks in *A* were detected with scipy.signal.find peaks. Writing *k*_*f*_ = ⌊*f N* ⌋ and

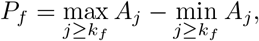

panel A and the protocol-check and additional panel-D points used *A*[*k*_0.6_ :] with prominence max(10^*−*7^, 0.08*P*_0.6_). Panel B and the corresponding panel-D points used *A*[*k*_0.5_ :] with prominence max(10^*−*5^, 0.1*P*_0.75_) and minimum peak distance 10. Focused panel-A rechecks used *A*[*k*_0.8_ :] within the current 701-point stage with prominence max(10^*−*7^, 0.08*P*_0.8_). Other find peaks arguments used SciPy defaults. Successive periods were calculated from consecutive peak times.

For Fig. 5A, preconditioning and post-switch integrations used BDF with

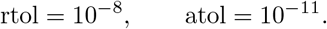

Post-switch trajectories were examined sequentially at

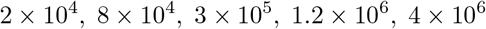

time units. A response was provisionally stationary when the maximum late-time amplitude over the four state variables was below 2 × 10^*−*5^, *d <* 3 × 10^*−*5^, and *r*_*x*_ *<* 10^*−*8^. Stationary states with *A <* 0.12 were assigned Low; responses approaching *S* = 2 were assigned Supermature only when the corresponding high-*S* branch was transversely stable; the remaining stationary responses were assigned Mature. A relaxed Low criterion required

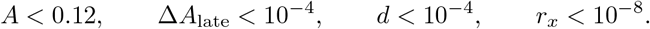

Oscillatory classification required at least four late-time peaks, late *A* amplitude above 10^*−*4^, period coefficient of variation below 0.15, *r*_*A*_ *>* 0.75, minimum *R >* 10^*−*12^, and an observation time exceeding both 10^5^ and five estimated periods.

Convergence toward the Disassembled state in the panel-A grid and associated checks, and in the protocol-check and additional points in panel D, was detected by the terminal event

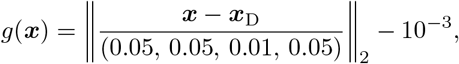

with decreasing-distance crossings only.

All Low-labeled panel-A grid points were subsequently screened against independently solved stable equilibria. Eighteen potential early-stop cases and 12 adjacent controls were re-integrated at tighter tolerances and extended observation times. For these focused rechecks, stationary classification required

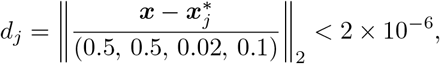

together with

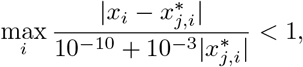

maximum late-time amplitude below 2 × 10^*−*6^, endpoint drift below 3 × 10^*−*6^, and *r*_*x*_ *<* 10^*−*10^. The focused oscillatory criterion used 0.9 *< r*_*A*_ *<* 1.1. Two early-stop Low assignments converged instead to the stable Mature equilibrium; the remaining 16 flagged cases and all 12 controls retained their original classifications.

For panel B and the panel-D points evaluated using the same procedure as panel B, stationary classification required a late *A* amplitude below 2 × 10^*−*5^, *a <* 3 × 10^*−*5^, and *r*_*x*_ *<* 10^*−*8^. Oscillatory classification required at least four sustained peaks, late *A* amplitude above 10^*−*4^, *r*_*A*_ *>* 0.75, period coefficient of variation below 0.25, and minimum late-time *R >* 10^*−*12^. For panel B, the final mode was evaluated over 3.6 × 10^5^ time units after return to *E* = 1. For panel D, observation times were extended near transitions; the response at *τ*_*F*_ = 11.9 remained nonstationary after 2 × 10^6^ time units and was retained as unresolved. Colored interval boundaries were placed between neighboring sampled *τ*_*F*_ values and were not treated as bifurcation locations.

For Fig. 5C, each preconditioning trajectory was integrated from the common Mature state to *T*_pre_ = 10^6^ using BDF with

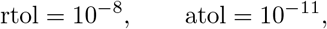

and dense output. The state at each sampled *T*_pre_ was used directly as the initial state after return to *E* = 1. Post-return trajectories were examined sequentially at

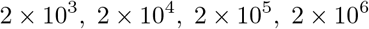

time units.

Classification was based on distances to independently solved stable Low and Mature equilibria,

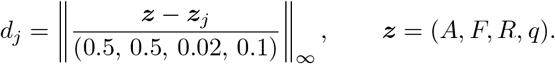

Mature classification required

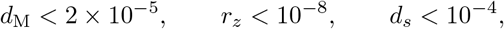

whereas Low classification required

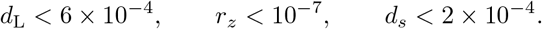

Responses satisfying neither criterion were retained as unresolved. All 5002 points in the final grid were ultimately classified as Mature or Low. Twelve representative and boundary-adjacent histories were recomputed with

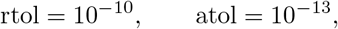

and observation times up to 4 × 10^6^; all retained the same classification.

For panels A and the corresponding panel-D checks, the terminal Disassembled event was evaluated before stationary and Oscillatory classification. For panel B and the panel-D points evaluated using the same procedure as panel B, stationary classification preceded the Oscillatory test. Panel C tested Mature, Low, and Disassembled classes only; unresolved cases were not forced into another mode.

### Independent verification and reproducibility

A periodic solution on the invariant boundary *R* = 0 has transverse reinforcement dynamics

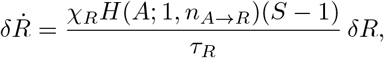

with Floquet multiplier

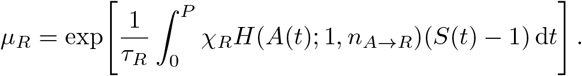

For the computed boundary cycle at *E* = 1 and *τ*_*F*_ = 12.5, *A*(*t*) *>* 0 and *S*(*t*) *>* 1 throughout the cycle, so *µ*_*R*_ *>* 1; the boundary cycle is therefore transversely unstable. The distinct positive-*R* periodic orbit at the same parameter values is orbitally stable, and the Mature equilibrium is locally stable, establishing Mature–Oscillatory coexistence from independent stability calculations.

The reproducibility packages retain the model implementation, numerical outputs, continuation or restart data where applicable, and the scripts used for the corresponding analyses and figures.

